# CK2α-dependent regulation of Wnt activity governs white matter development and repair

**DOI:** 10.1101/2023.04.11.536369

**Authors:** Chih-Yen Wang, Zhongyuan Zuo, Kyoung In Kim, Hugo J. Bellen, Hyun Kyoung Lee

## Abstract

Wnt signaling plays an essential role in developmental and regenerative myelination in the CNS. The Wnt signaling pathway is comprised of multiple regulatory layers; thus, how these processes are coordinated to orchestrate oligodendrocyte development remains unclear. Here we show CK2α, a Wnt/β-catenin signaling Ser/Thr kinase, phosphorylates Daam2, inhibiting its function and Wnt-activity during oligodendrocyte development. Intriguingly, we found Daam2 phosphorylation differentially impacts distinct stages of oligodendrocyte development, accelerating early differentiation followed by decelerating maturation and myelination. Application towards white matter injury revealed CK2α-mediated Daam2 phosphorylation plays a protective role for developmental and behavioral recovery after neonatal hypoxia, while promoting myelin repair following adult demyelination. Together, our findings identify a novel regulatory node in the Wnt pathway that regulates oligodendrocyte development via protein phosphorylation-induced signaling complex instability and highlights a new biological mechanism for myelin restoration.

**Significance:** Wnt signaling plays a vital role in OL development and has been implicated as an adverse event for myelin repair after white matter injury. Emerging studies have shed light on multi-modal roles of Wnt effectors in the OL lineage, but the underlying molecular mechanisms and modifiable targets in OL remyelination remain unclear. Using genetic mouse development and injury model systems, we delineate a novel stage-specific function of Daam2 in Wnt signaling and OL development via a S704/T7-5 phosphorylation mechanism, and determine a new role of the kinase CK2α in contributing to OL development. In-depth understanding of CK2α-Daam2 pathway regulation will allow us to precisely modulate its activity in conjunction with Wnt signaling and harness its biology for white matter pathology.

## Introduction

Oligodendrocytes (OLs) are myelin-producing cells that provide functional and metabolic support to axons in the central nervous system (CNS) (1). In white matter disorders, such as hypoxic ischemic encephalopathy (HIE) and multiple sclerosis (MS), loss of OLs and myelin sheaths are early hallmarks of disease initiation and progression (2, 3). OL precursor cells (OPCs) are recruited to the lesion for tissue repair, but secretory molecules from the adjacent tissue block OL differentiation, thereby limiting remyelination (4-7). Importantly, upregulation of canonical Wnt signaling has been reported in MS patients (8, 9), and aberrant activation of Wnt signaling is well accepted as an adverse event for remyelination. However, manipulating Wnt regulators using genetic models has produced inconsistent outcomes (10, 11), possibly because these Wnt components interact with other pathways, affect transcriptional partners at different stages of OL lineage, or have Wnt-independent functions (7, 10, 12). Therefore, it is imperative to delineate the temporal dynamics and molecular mechanism of Wnt signaling at different stages of OPCs/OLs.

Cytoskeletal remodeling is crucial for stage-specific OL lineage progression. While actin polymerization is required for OL process extension during early differentiation and ensheathment stages, actin depolymerization is necessary for myelin wrapping at the maturation period (13). Formin proteins play a key role in cellular morphogenesis by mediating actin assembly and cytoskeletal remodeling (14). Daam2 (Dishevelled-associated activator of morphogenesis 2) is a formin member that is a positive regulator of canonical Wnt signaling during embryonic spinal cord patterning (15). Daam2 overexpression greatly inhibits OL differentiation, where loss of Daam2 promotes differentiation during development and after white matter injury (16, 17). Consistent with this finding, Daam2 is upregulated in demyelinated lesions in conjunction with a higher Wnt tone in HIE and MS patients (11, 16, 17). However, we recently found that loss of Daam2 leads to an abnormal myelin formation which returns to normal at an older stage (18). Hence, these studies raise two important questions. 1) Does Wnt signaling play a negative role in early differentiation and a positive role during maturation and myelination? 2) What are the molecular mechanisms that regulate the activity of Daam2?

CK2, a serine/threonine kinase, interacts with multiple Wnt components and positively regulates Wnt activity (19-21). While it is established that CK2 subunits are essential for brain development and OPC production (22, 23), their potential roles and targets in OL differentiation via Wnt signaling remain to be determined. In this study, we discovered that phosphorylated Daam2 at S704/T705 attenuates Wnt/β-catenin signaling in OL lineage and promotes OL differentiation but subsequently decelerates myelination. We identified CK2α as the kinase that phosphorylates Daam2 followed by weakening Wnt signaling complex. Moreover, in white matter injury models, both CK2α overexpression and Daam2 phosphorylation were found to promote tissue repair. Our findings establish a novel role for CK2α in blunting the adverse effect of Daam2-mediated Wnt signaling for OL differentiation, suggesting a new regulatory pathway for white matter diseases.

## Results

### Daam2 Phospho-mimetic mutant promotes OL differentiation

To identify possible regulators of Daam2, we performed Daam2 immunoprecipitations (IP) on mouse cortical tissues followed by mass spectrometric (IP-MS) analysis. We found that residues S704 and T705 in the FH2 domain of Daam2 are phosphorylated. The amino acid sequence is highly conserved among species in Daam2 orthologues (**Fig. 1A**). The S704 phosphorylation (S656 in humans) of Daam2 has been previously reported (24, 25). To investigate the effect of phosphorylation, we introduced phospho-null (S704A/T705A, denoted as A-mutant), or phospho-mimetic (S704E/T705E, denoted as E-mutant) mutants into Daam2, and transfected them into primary OPCs (**Fig. 1A-1B**). We confirmed that wild-type (WT) Daam2 suppresses OL differentiation as evidenced by a reduced number of mature OL markers including MAG^+^ and MBP^+^ cells after 2 and 4 days of OL differentiation, respectively (***SI Appendix,* Fig. S1A-C**). The overexpression of the A-mutant had a comparable negative effect on OL differentiation whereas the E-mutant increased the number of MAG^+^ and MBP^+^ cells with morphological complexity (**Fig. 1B**; ***SI Appendix,* Fig. S1D-F**).

**Fig. 1.**
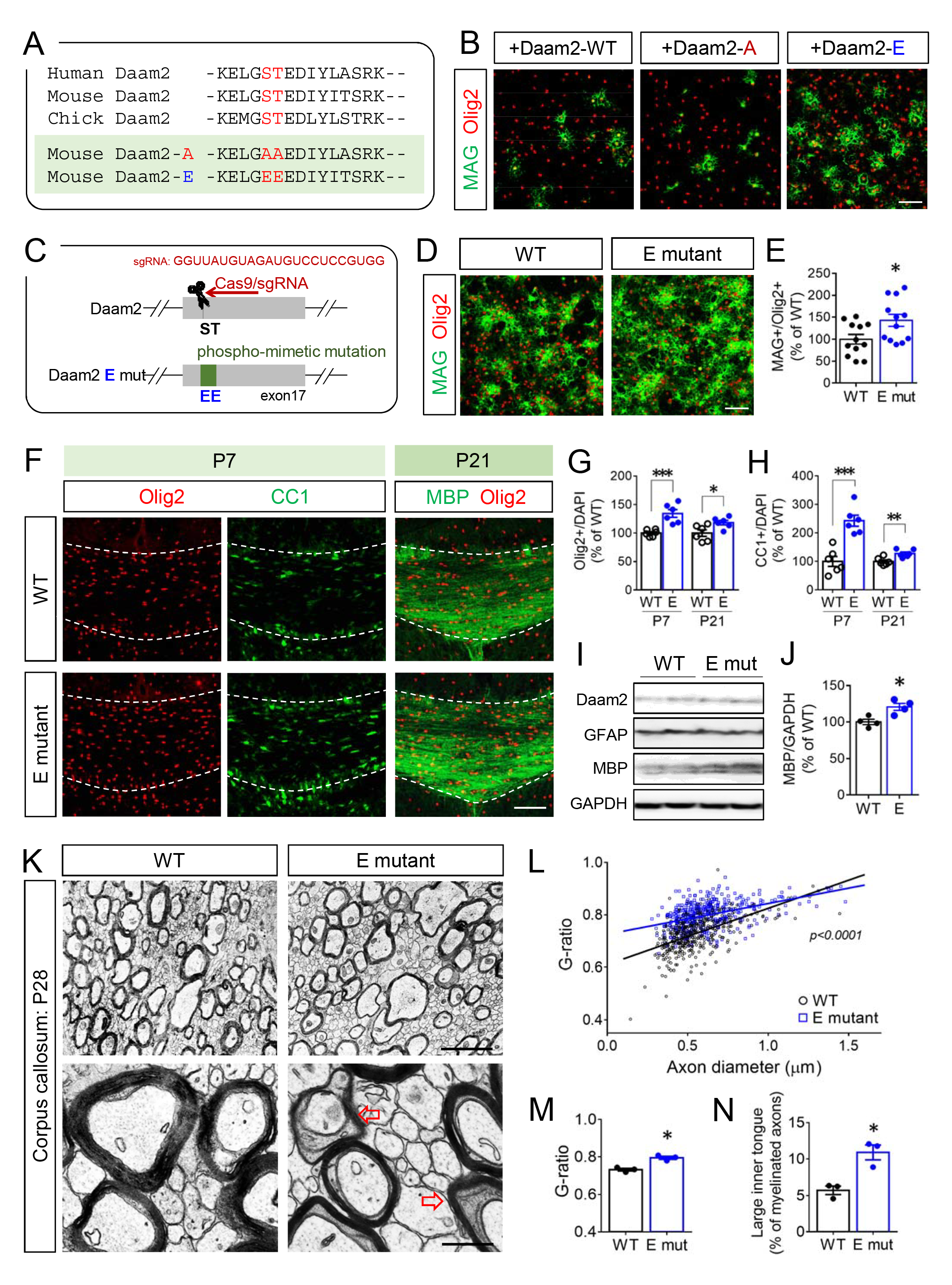
Phospho-mimetic mutation of Daam2 accelerates OL differentiation but delays myelination. (A) Phosphorylation motif of Daam2 in mouse, human and chick are shown. For following experiments, phospho-mimetic (E-mut) and phospho-null mutation (A-mut) of Daam2 were used. (B) Daam2-WT, E-mut, and A-mut were transfected into primary OPCs followed by differentiation for 2 days. (C) E-mutant mice were produced with CRISPR-Cas9-based technique. The guide RNA sequence for the E-mutation is shown. (D) *In vitro* OL differentiation was assessed using WT and the E-mutant OL culture. (E) The number of MAG^+^ cells for 2-day differentiation was quantified. (F-H) P7 and P21 brains from WT and the E-mutant mice were analyzed by immunofluorescence. Olig2^+^ and CC1^+^ (APC^+^) cells in the corpus callosum were counted. (I-J) P21 cortical tissues containing the corpus callosum from WT and the E-mutant were analyzed by western blot. The MBP protein levels were quantified. (K) The myelin structure in the corpus callosum from WT and the E-mutant mice at P28 were subjected to electron microscopy. (L-M) Axon diameter and g-ratio from each myelinated axon were measured. (N) Axons with enlarged inner tongue microstructure (red arrows in K) were also counted. The data were collected from at least 3 independent experiments or animals for each group. Data were presented as mean ± SEM and normalized to WT. *P < 0.05, **P < 0.01, ***P < 0.001 versus WT. Scale bar, 100 μm in B, D, F; 2 μm (upper) and 0.5 μm (lower) in K.

To examine the endogenous function of Daam2 phosphorylation *in vivo*, we generated knock-in mice bearing the Daam2 E-mutation in the endogenous locus (**Fig. 1C**). In accordance with our *in vitro* findings, OPCs from the E-mutant brain produced more MAG^+^ and MBP^+^ OLs than WT brains (**Fig. 1D-1E**; ***SI Appendix,* Fig. S1G**). During the myelinogenesis period in the brain, we also found more Olig2^+^ OL lineage cells and CC1^+^ mature OLs in the E-mutant corpus callosum at P7 and P21 than in the WT (**Fig. 1F-1H**; ***SI Appendix,* Fig. S1J**). *In vitro* data indicate that E-mutation did not affect OPC proliferation (***SI Appendix,* Fig. S1H**), but facilitates OPC specification and OL lineage progression from neural stem cells (NSCs; ***SI Appendix,* Fig. S1I**). In addition, MBP levels were upregulated in the E-mutant brain (**Fig. 1I-1J; *SI Appendix,* Fig. S1K**), yet the thickness of the myelin sheath in the corpus callosum was reduced (**Fig. 1K-1M**). We also observed more axons surrounded by enlarged inner tongue structures, the cytoplasmic space between the axons and the myelin, in the E-mutant corpus callosum at P28 (**Fig. 1K and 1N**; **arrows in 1K**). However, myelin integrity in the E-mutant was restored in adulthood (***SI Appendix,* Fig. S1L**), suggesting it takes a longer time to complete axon ensheathment in the E-mutant brain. Yet, we did not observe significant differences between the E-mutant and WT brains with respect to astrocytes, another group of Daam2-expressing cells in the CNS (***SI Appendix,* Fig. S1M-O**). Our results indicate that the S704/T705 phosphorylation could alter Daam2 functions in the OL lineage progression.

### Dynamic transcriptomic remodeling in early and late OLs by Daam2 phosphorylation

To identify the molecular signatures that underlie the observed pro-differentiation effects of the Daam2 E-mutant, we performed transcriptome analysis using 10X Genomics single cell-RNA sequencing (scRNA-seq) in P21 brains. After unbiased clustering, four clusters of OL lineage cells (*Pdgfra*^+^ OPCs, *Cspg4*^+^ OPCs, early differentiated OLs, late differentiated OLs) were identified using specific OL stage markers such as *Pdgfra*, *Cspg4*, *LncOL1* and *Mobp* (**Fig. 2A-2B**) from WT and E-mutant brains. Specifically, early and late gene signatures for OL progression were identified. For the early genes (those correlated with the early OL marker *LncOL1*), *Cemip2, Itpr2, and Gpr17* were significantly upregulated by the E-mutation in early OLs compared to WT (**Fig. 2C**). In contrast, the intermediate gene *Ctps* showed a reversed trend in the early OL cluster (**Fig. 2D**). Notably, the late genes, *Aspa*, *Car2* and *Ptgds* (as mature OL markers) were largely downregulated by the E-mutation in both early and late OL clusters (**Fig. 2D**). Collectively, these transcriptomic changes are consistent with our observation that Daam2 phosphorylation accelerates OL differentiation but decelerates maturation and myelination (**Fig. 1**).

**Fig. 2.**
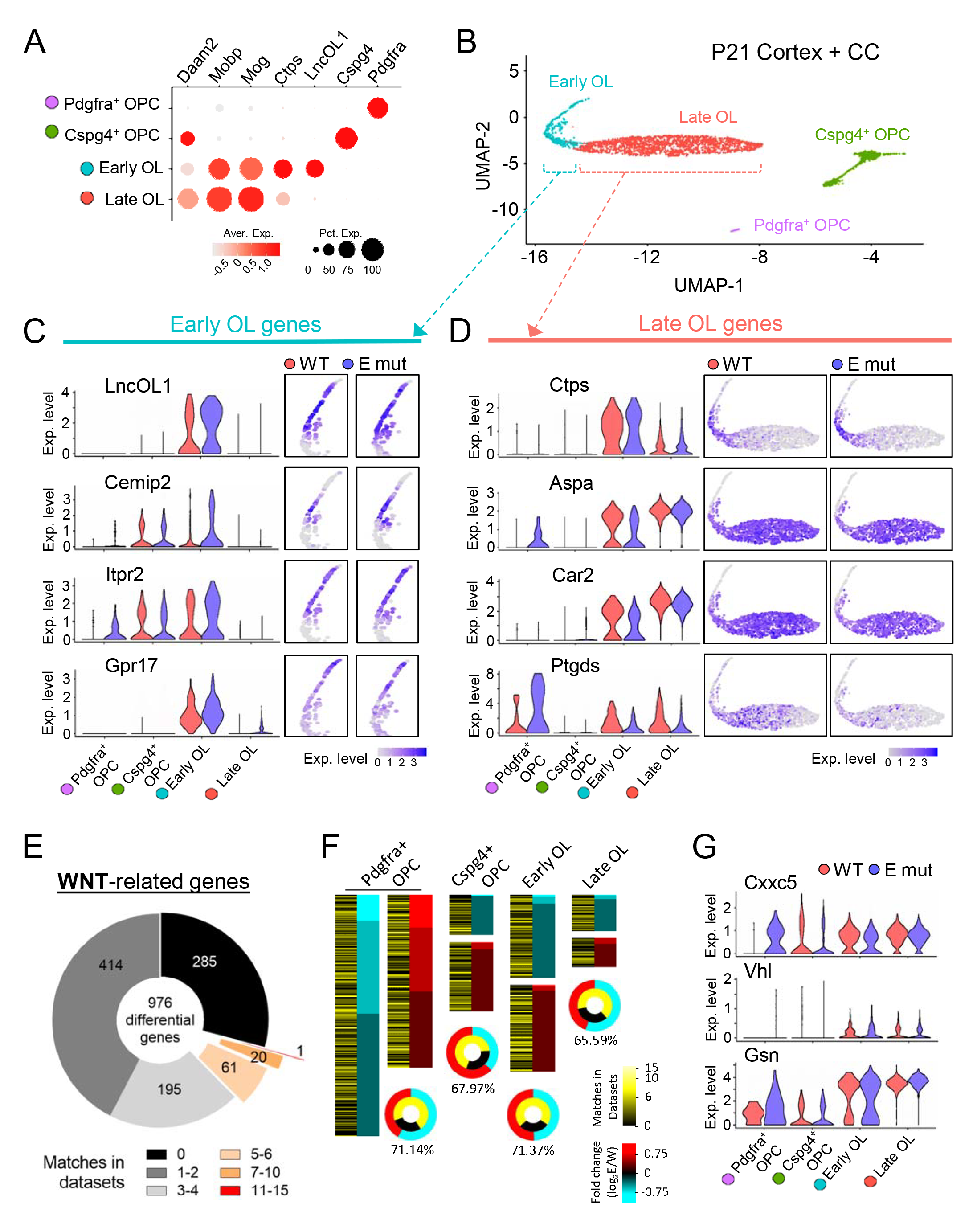
Early genes are upregulated but late genes are downregulated in the E-mutant OLs by scRNA-seq analysis. (A) The cell identities of clusters from scRNA-seq using in P21 brains were determined by specific markers, *Pdgfra* (OPCs), *Cspg4* (OPCs), *LncOL1*, *Ctps* (early OLs), *Mog*, and *Mobp* (late OLs). (B) OPC/OL clusters were visualized by dimension reduction plot, UMAP. (C-D) Early OL genes (*LncOL1*, *Cemip2*, *Itpr2*, and *Gpr17*), intermediate gene (*Ctps*), late/maturation genes (*Aspa*, *Car2*, and *Ptgds*) of WT and the E-mutant are shown by violin plots (left panels). The locations of cells expressing the genes were displayed in UMAP of WT and the E-mutant (right panels). (E) 976 differentially expressed genes (DEGs; Log2-Fold change > 0.25 and P value < 0.05) between WT and the E-mutant were aligned with 28 datasets containing Wnt-related gene list (also see Table S1). The number of genes that matched with Wnt datasets are shown in different divisions in the pie chart. (F) For 4 clusters, genes upregulated in E-mutant are marked in red, and downregulated in cyan. Genes that match Wnt datasets are labeled in dark to bright yellow (low to high matches). (G) Wnt-related genes, Cxxc5 (Wnt inhibitor gene), Vhl and Gelsolin (also from Daam2 studies (17, 18)) are shown in violin plots.

To explore how Daam2 phosphorylation regulates OL lineage progression, differentially expressed genes (DEGs) between WT and the E-mutant were subjected to GO and KEGG enrichment analysis. DEGs downregulated in the E-mutant OPCs/OLs are enriched for ER-mediated protein processing and lipid/cholesterol metabolism (***SI Appendix,* Fig. S2**). In contrast, DEGs upregulated by the E-mutation are involved in multiple signaling processes, including the Wnt pathway. Since Daam2 is a positive modulator of canonical Wnt signaling, we further examined whether these DEGs result from alteration of Wnt signaling. Crosschecking with 28 published datasets containing Wnt-related genes (**Table S1**) revealed that over 70% of the DEGs are associated with Wnt signaling (**Fig. 2E**). We also observed higher portions of Wnt-related DEGs in *Pdgfra*^+^ OPCs and early OLs than in late OLs (**Fig. 2F**). For example, *Cxxc5*, a Wnt negative feedback regulator and a transcriptional activator for myelin genes, was elevated in *Pdgfra*^+^ OPCs but reduced in early OLs, suggesting a temporally-specific regulatory mechanism of Wnt signaling during OL differentiation (**Fig. 2G**). Moreover, we also validated upregulation of *Vhl* and *Gsn* from Wnt-regulated DEGs in the E-mutant OLs (**Fig. 2G**), previously found to promote OL differentiation (17, 18). Together, these results suggest that phosphorylation of Daam2 regulates Wnt signaling in the entire OL lineage and controls OL differentiation and maturation respectively.

### Daam2 regulates Wnt signaling in the OL lineage in a stage-specific manner

We next investigated whether Daam2 phosphorylation regulates Wnt activity in the OL lineage in a stage-specific manner. To determine endogenous Wnt activity in primary OLs, we assessed the nuclear β-catenin level at differentiation day 2 (early; **Fig. 3A**) and day 4 (late; **Fig. 3E**). Interestingly, early OLs showed a large amount of cytosolic β-catenin (***SI Appendix,* Fig. S3A**), while more β-catenin accumulated in the nucleus in late OLs (***SI Appendix,* Fig. S3B**). To further induce canonical Wnt signaling, we treated early and late OLs with canonical Wnt ligands, Wnt3a and Wnt7a, which caused a two-fold increase in nuclear β-catenin levels (**Fig. 3B-3C and 3F-3G; *SI Appendix,* lane 5 vs 6 in Fig. S3A-S3B**). In contrast, nuclear β-catenin levels remained unchanged in E-mutant OLs after Wnt3a/7a treatment, indicating that Daam2 phosphorylation severely attenuated ligand-based Wnt/β-catenin signaling (***SI Appendix,* Fig. S3A-S3B**; **lane 7 vs 8**). To further understand the relationship between Wnt activity and OL differentiation, we examined early differentiation versus late/terminal differentiation after Wnt3a/7a treatment. Wnt3a/7a greatly reduced early differentiation, with less MAG expression in the WT than in the E-mutant (**Fig. 3B and 3D**). Wnt3a/7a treatment also improved MBP levels and membrane structures at a later stage of OL differentiation in the WT (**Fig. 3F and 3H; *SI Appendix,* lane 1 vs 2 in Fig. S3B**), while E-mutant OLs were insensitive to Wnt ligands treatment (**lane 3 vs 4**).

**Fig. 3.**
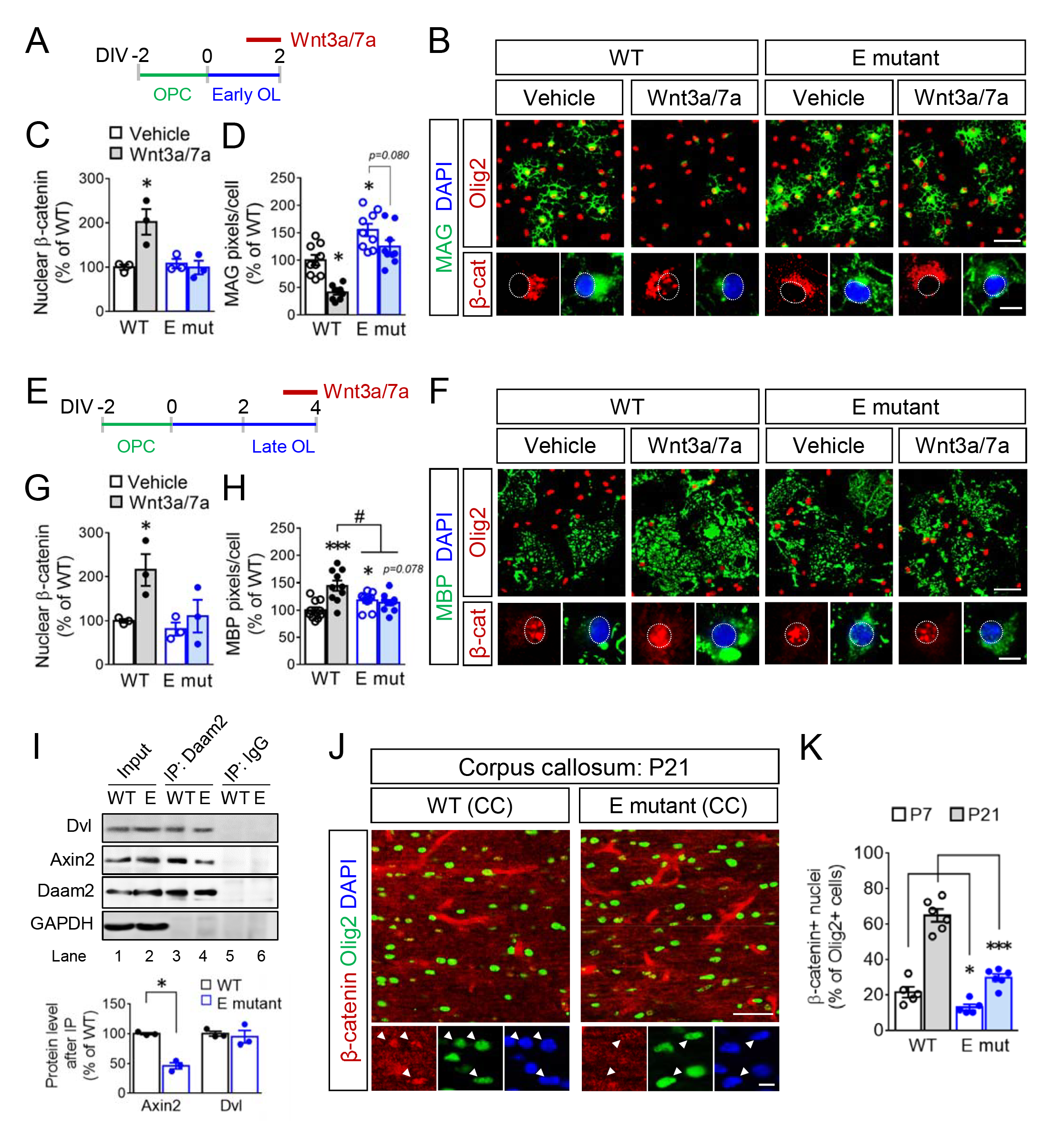
Exogenous Wnt/β-catenin signaling were blocked in the E-mutant OLs. (A) Wnt3a/7a were treated to early OLs of WT and the E-mutant for 24 hours prior to harvesting. (B) *In vitro* differentiation by MAG expression in early OLs after Wnt3a/7a treatment were assessed by immunofluorescence. The subcellular locations of β-catenin were also visualized. (C) β-catenin levels in the nucleus fractions of early OLs were analyzed and quantified by western blot (also see Figure S4A). (D) The MAG expression in B was quantified. (E-H), Similar to A-D, Wnt3a/7a were treated to late OLs of WT and the E-mutant. The nuclear β-catenin (G) and MBP expression (H) in late OLs were measured. (I) Daam2 IP was conducted using WT and the E-mutant brain at P21, Wnt complex components, Dvl and Axin2 were blotted and the protein enrichment was quantified in the lower panel. (J-K), β-catenin distributions were evaluated by immunofluorescence in the WT and E-mutant corpus callosum at P7 and P21. Nuclear β-catenin was identified by overlapping with DAPI^+^ nuclei and by showing higher intensity than the surroundings. OL lineage cells and mature OLs were labeled by Olig2 and CC1, respectively. The data were collected from at least 3 independent experiments or animals for each group. Data were presented as mean ± SEM and normalized to WT/vehicle. *P < 0.05, **P < 0.01, ***P < 0.001 versus WT/vehicle; ^#^P < 0.05, ^##^P < 0.01 versus WT/Wnt. Scale bar, 50 μm (upper), 10 μm (lower) in B, F, J.

Given that Daam2 is required for Wnt signalosome complex formation, we investigated whether Daam2 phosphorylation interferes its interaction with Wnt components. Although the interaction of Daam2 and Dishevelled (Dvl) was not altered by the E-mutation in the brain, there was a reduction in the association between Axin2 and the Daam2 E-mutant (**Fig. 3I**). This evidence explains that Daam2-Axin2 interaction is important for Daam2 phosphorylation-mediated Wnt signaling cascade. To validate our *in vitro* findings, we assessed the β-catenin protein expression pattern in WT and the E-mutant corpus callosum at P7 and P21 (**Fig. 3J**). At P7, nuclear β-catenin was observed in ∼20% of Olig2^+^ population of cells in WT which increased to ∼70% at P21 (**Fig. 3J-K**). In comparison, the overall levels of nuclear β-catenin were downregulated in the E-mutant mice. Together, these observations demonstrate dynamic changes in the machinery and function of Wnt/β-catenin signaling in early versus late OLs, and this pathway is affected by Daam2 phosphorylation.

### CK2***α***-mediated Daam2 phosphorylation stimulates OL differentiation

To identify the kinases responsible for S704/T705 phosphorylation, we conducted motif analysis by using the NetPhos3.1 database (26). We found CK2 among the candidates from IP-MS data (***SI Appendix,* Fig. S4A-S4B**), and we confirmed that its major catalytic subunit, CK2α, interacts with Daam2 in primary OLs (***SI Appendix,* Fig. S4C**). To investigate whether CK2α directly phosphorylates Daam2, we performed *in vitro* kinase assays using either purified Daam2 proteins or synthesized peptides containing the phosphorylation site (**Fig. 4A**). CK2α phosphorylates both Daam2 and the synthesized peptides, and both the S704A or T705A mutation blocked this phosphorylation (**Fig. 4A-4B**). We further examined whether CK2α phosphorylates Daam2 in primary OLs and found that CK2α overexpression increased phospho-serine (p-Ser) levels of WT-Daam2, but not of the A-mutant protein levels (***SI Appendix,* Fig. S4D**). Moreover, CK2α and Daam2 proteins were sequentially upregulated in the cytosolic compartment concomitant with OL lineage progression (**Fig. 4C**). These results suggest a regulatory pathway of CK2α-Daam2 in OLs.

**Fig. 4.**
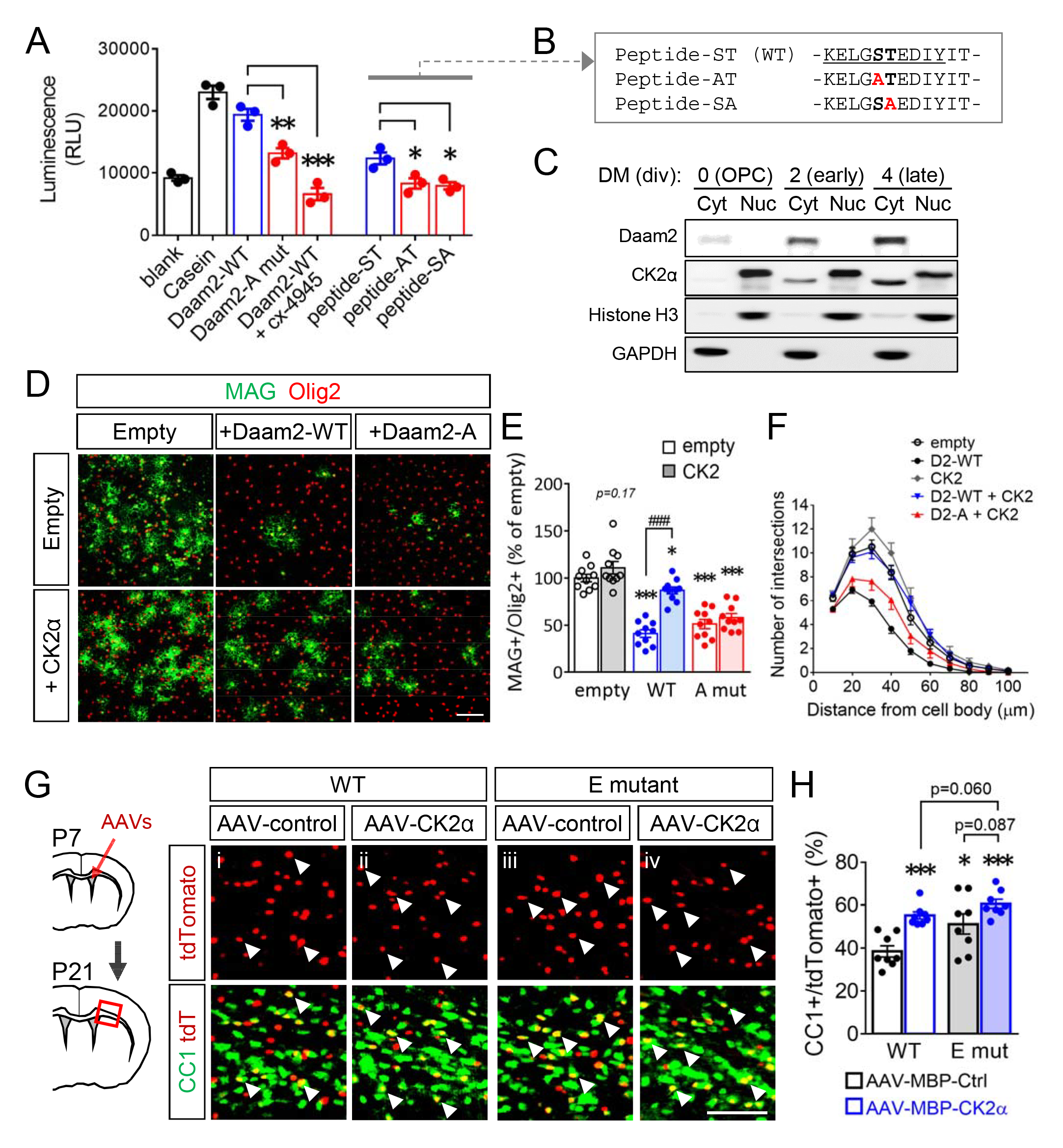
CK2α phosphorylates Daam2 and facilitates OL differentiation. (A-B) Purified Daam2 proteins and synthesized peptides (sequences shown in B) were subjected to *in vitro* CK2 kinase assay. Blank: no substrate; Casein: positive control. (C) Endogenous CK2α expression was blotted in nucleus and cytosol fractions from different stages of OPC/OL culture. (D) *In vitro* differentiation was evaluated by immunofluorescence after transducing Daam2 and CK2α. (E) The number of MAG^+^ cells/Olig2^+^ cells were counted. (F) The number of processes extended from a cell body at different distances was analyzed by sholl analysis. (G-H) AAV-MBP-CK2α and AAV-MBP-control were injected intracerebroventricularly into P7 pups respectively. CC1^+^ cells with tdtomato reporter were visualized by immunofluorescence. % of injected cells that became mature OLs was quantified as CC1/tdtomato. Data from at least 3 independent experiments or animals for each group were presented as mean ± SEM and normalized to empty or WT/AAV-control. *P < 0.05, **P < 0.01, ***P < 0.001 versus WT proteins in A, versus empty in E, versus WT/AAV-control in H; ^###^P < 0.001 versus Daam2-WT. Scale bar, 100 μm in D, 50 μm in G.

Next, we tested whether CK2α facilitates OL differentiation via Daam2 phosphorylation. Although CK2α overexpression showed an increasing trend in OL differentiation *in vitro*, CK2α reversed the inhibitory effect of Daam2 (**Fig. 4D-4F**). In agreement with previous data, OL differentiation was also reduced by CK2 inhibitors, cx-4945 and DMAT (***SI Appendix,* Fig. S5A-S5B**) or a CK2α dominant negative mutant (K68A; dnCK2α; ***SI Appendix,* Fig. S5C-S5D**), indicating a requisite role of CK2 kinase function. To examine the function of CK2α in mice *in vivo*, we created an adeno-associated viruses (AAVs) carrying the CK2α gene as well as the tdtomato reporter under the control of the MBP promoter to express the gene selectively in differentiating OLs. The virus was injected intracerebroventricularly into mouse brains at P7 (***SI Appendix,* Fig. S5E**). The upregulation of CK2α by AAV-CK2α was confirmed in tdtomato^+^ cells when compared to the AAV-controls using the WT brain (***SI Appendix,* Fig. S5F**). Additionally, most of the tdtomato^+^ cells were restricted to the corpus callosum at P21 as Olig2^+^ OL lineage (***SI Appendix,* Fig. S5G-S5H**). As a result, OL-specific overexpression of CK2α increased the abundance of CC1^+^ cells among tdtomato^+^ cells (**Fig. 4G-4H; panel i vs ii**), confirming the stimulatory role of CK2α in OL differentiation *in vivo*.

To further validate the CK2α-Daam2 pathway *in vivo*, we also introduced the AAVs into the E-mutant brains. As the E-mutant mice displayed early OL differentiation, there were more CC1^+^/tdtomato^+^ cells in the E-mutant brains receiving the AAV-control as compared to the WT brains (**Fig. 4G-4H; panel i vs iii**). However, the E-mutant brains receiving AAV-CK2α did not show further enhancement by CK2α overexpression and by the E-mutation, respectively (**Fig. 4G-4H; panel iv vs ii, iii**), confirming CK2α and Daam2 phosphorylation share the same pathway in this model. Therefore, these data provide compelling evidence that CK2α promotes OL differentiation by phosphorylating Daam2.

### Daam2 phosphorylation improves functional recovery after neonatal hypoxic injury

Given that Wnt signaling and are upregulated in white matter lesions of human HIE (16), we next evaluated whether attenuating Daam2 function by modulating its phosphorylation state improves recovery in white matter injury models. In a mouse model of neonatal injury that mimics human HIE (16, 27), brain development was impaired in a hypoxic environment (10.5% O_2_) from P3 to P11 (**Fig. 5A-5B**). OL differentiation was significantly impaired as CC1^+^ OLs were barely detected in the corpus callosum immediately after hypoxic injury at P11 (**Fig. 5C**) in conjunction with Daam2 upregulation (***SI Appendix,* Fig. S6A**). Notably, hypoxia also reduced the number of PDGFRα^+^ OPCs (**Fig. 5D**). In comparison with Daam2 WT controls, there were more CC1^+^ mature OLs and PDGFRα^+^ OPCs in the E-mutant corpus callosum after injury (**Fig. 5C-5D**). Moreover, after a 7-day recovery period post-injury, more CC1^+^ OLs were generated in the E-mutant brain compared to WT (**Fig. 5E-5F**). MBP immunoreactivity at the cingulum, an early myelination event, was also higher in the E-mutant mice than in WT (**Fig. 5G**). Moreover, we observed significantly more myelinated axons and slightly increased myelin sheath thickness in the E-mutant corpus callosum compared to WT (**Fig. 5H-5K**). There was no difference in the development of neurons and astrocytes between WT and E-mutant brains after injury at P18 (***SI Appendix,* Fig. S6C-S6D**).

**Figure 5.**
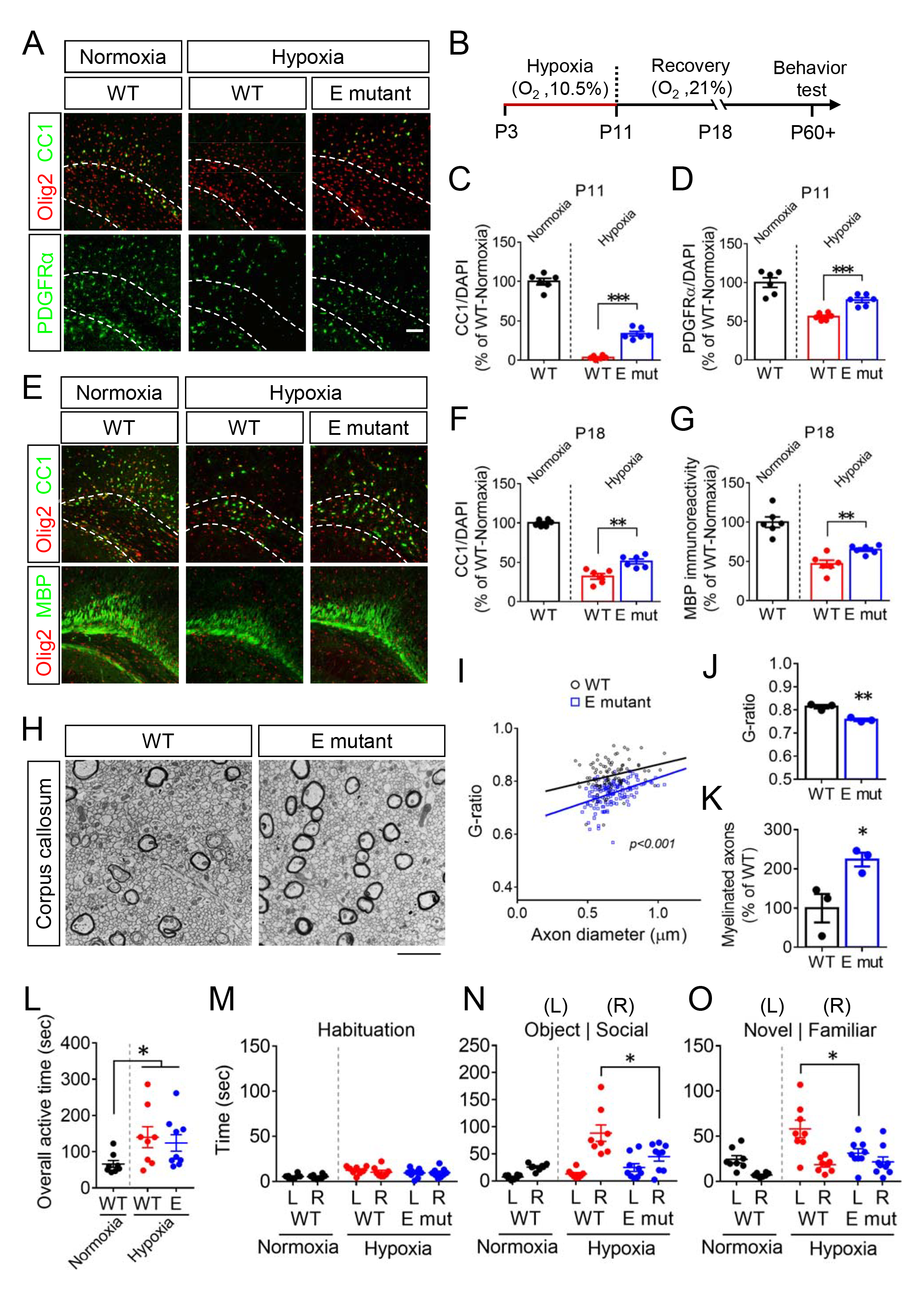
The E-mutant mice show better functional recovery after postnatal hypoxic injury. The brains of P11 (A-D) and P18 (E-G) WT and the E-mutant mice, that suffered from postnatal hypoxic injury (from P3 to P11), were analyzed by immunofluorescence. The numbers of Olig2^+^, CC1^+^, and PDGFRα^+^ cells were counted within the corpus callosum (dash line), and the immunoreactivity of MBP was measured. (H) The myelin structure in the corpus callosum from WT and the E-mutant mice at P18 after postnatal hypoxic injury were subjected to electron microscopy. (I-J) Axon diameter and g-ratio from each myelinated axon were measured. (K) The number of myelinated axons was also counted. (L-O) The social behaviors of 2-month-old mice were evaluated by 3-chamber sociability test, including overall active time during tests (L), habituation (M), preference to an object or a mouse (N), and preference for a new stranger mouse or a familiar one (O). Data from at least 3 (H-K) or 6 animals for each group were presented as mean ± SEM. *P < 0.05, **P < 0.01, ***P < 0.001. Scale bar, 100 μm in A, E; 2 μm in H.

White matter development and neural connectivity are closely related to social behaviors (28). Both human HIE patients and rodents with neonatal hypoxia display functional deficits in childhood and show long-term effects in adolescence and adulthood, such as ADHD-like behaviors and social problem (27, 29, 30). To investigate whether Daam2 phosphorylation is critical for social behavior development after neonatal hypoxic injury, mice were challenged under 10.5% O_2_ hypoxic condition from P3 to P11, and then subjected to a three-chamber sociability test at two months of age (**Fig. 5B**). Similar to previous findings, mice with neonatal hypoxic injury were more active than normal mice (**Fig. 5H**), despite that there is no behavioral difference between WT and E-mutant mice during habituation (**Fig. 5I**). Neonatal hypoxia WT mice showed a higher preference for a stranger mouse over a novel object (**Fig. 5J**), and spent more time with a stranger mouse than with a familiar one (**Fig. 5K**). In contrast, the Daam2 E-mutation mitigated abnormal social preferences induced by hypoxia injury (**Fig. 5J-5K**). These data suggest a protective role of Daam2 phosphorylation for developmental and behavioral functional recovery after neonatal hypoxic injury.

### CK2***α***-induced Daam2 phosphorylation facilitates myelin repair after lysolecithin-induced demyelination

We previously discovered that Daam2 is expressed in OPCs in human MS lesions (17). To examine the function of Daam2 phosphorylation via CK2α in a mouse model of demyelination that mimics MS (31), we injected 2% lysolecithin with AAV-MBP-CK2α and -control virus in the corpus callosum of E-mutant and WT brains, respectively (**Fig. 6A**). After 14 days post-injury (dpi), we confirmed upregulation of Daam2 (***SI Appendix,* Fig. S6B**) and evaluated remyelination surrounding the lesion. OL-specific CK2α overexpression significantly increased the number of *Mbp*^+^ and *Plp*^+^ cells around the lesions compared to the control (**Fig. 6A-6C; panel i vs ii**), with no difference in *Pdgfra*^+^ cell numbers. Specifically, there were more CC1^+^ OLs among tdtomato^+^ cells infected with AAV-MBP-CK2α than those with AAV-control (**Fig. 6E-6G; panel i vs ii**). On the other hand, there were more *Mbp*^+^ and *Plp*^+^ OLs for tissue repair in the E-mutant brain compared to the WT brain (**Fig. 6A-6C; panel i vs iii**). The tdtomato^+^ cells infected with AAV-control generated a higher portion of CC1^+^ OLs in the E-mutant brain than in the WT brain (**Fig. 6E-6G; panel i vs iii**). Similar to our observations in developing brains (**Fig. 4G-4H**), CK2α overexpression in the E-mutant OLs did not show additional or synergistic effect for white matter repair as compared to that in WT OLs in the lesions (**Fig. 6E-6G; panel ii vs iv**). Moreover, we observed more myelinated axons as well as thicker myelin sheaths in the corpus callosum with CK2α overexpression and/or E-mutation (**Fig. 6H-6J)**. There is no difference in reactive astrocytes between E-mutant and WT brains at 14 dpi (***SI Appendix,* Fig. S6E-S6G**). These findings demonstrate that the CK2α-Daam2 pathway facilitates remyelination after white matter injury.

**Fig. 6.**
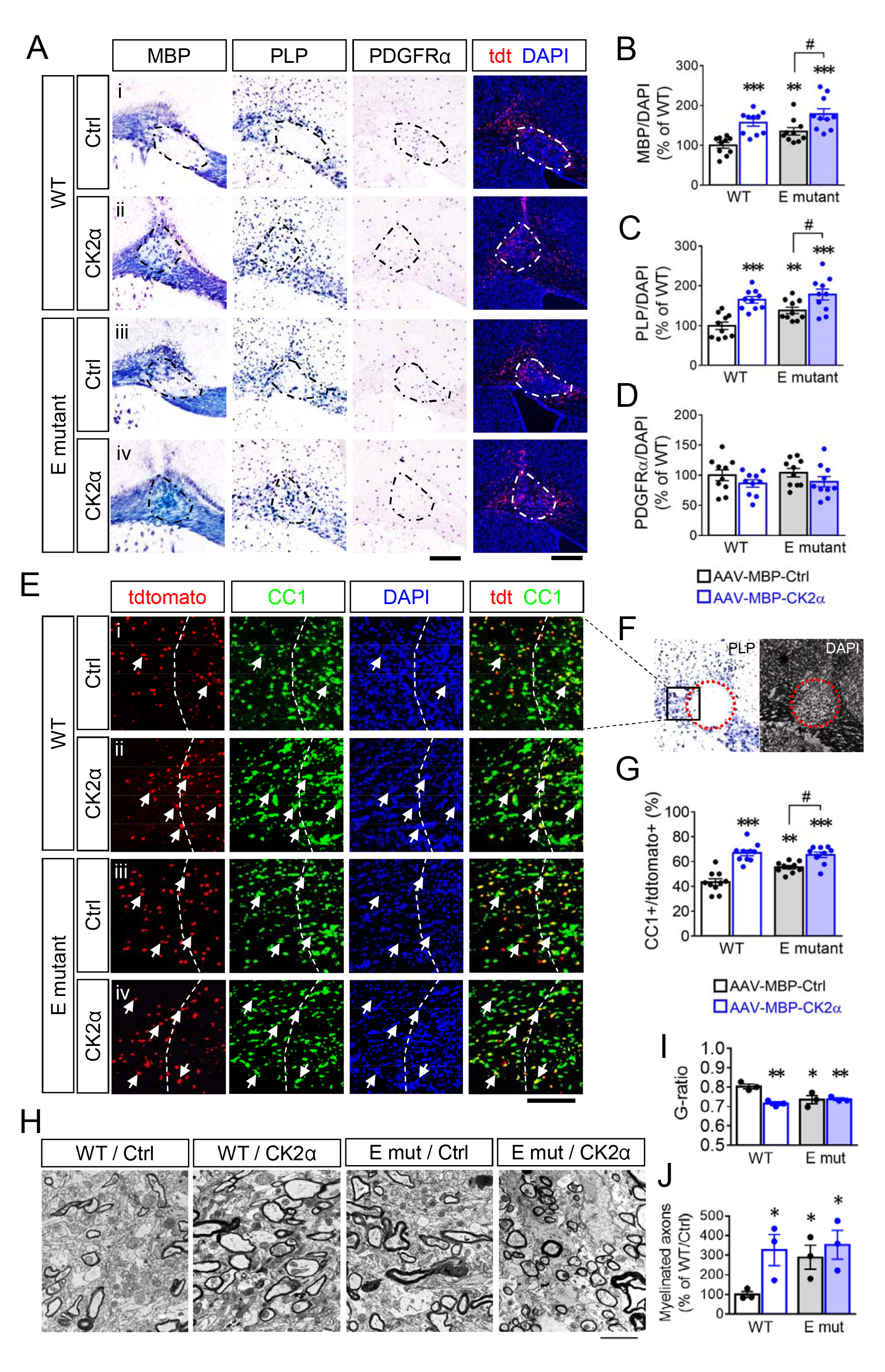
The E-mutant mice show improved tissue repair after lysolecithin-induced demyelination. At 14 days after injecting lysolecithin and AAVs into corpus callosum in adult mice, brain tissues containing the lesions were assessed by in situ hybridization for *Mbp*, *Plp* and *Pdgfr*α (A-D). The lesions were visualized by tdtomato reporter and accumulative DAPI. The number of *Mbp*^+^, *Plp*^+^, *Pdgfr*α^+^ cells, and DAPI in the lesion (dash line) area were counted. (E-G) The lesioned areas were also analyzed by immunofluorescence for CC1^+^ and tdtomato^+^ infected cells. % of injected cells that became mature OLs was quantified as CC1/tdtomato. (H) The myelin structure in the corpus callosum from WT and E-mutant mice at 14 days after receiving lysolecithin and AAVs were subjected to electron microscopy. (I) G-ratio from each myelinated axon was measured. (J) The number of myelinated axons was also counted. Data from at least 10 animals for each group were presented as mean ± SEM. **P < 0.01, ***P < 0.001 versus WT/AAV-control, ^#^P < 0.05 versus E-mut/AAV-control. Scale bar, 200 μm in A; 100 μm in E; 2 μm in H.

**Fig. 7.**
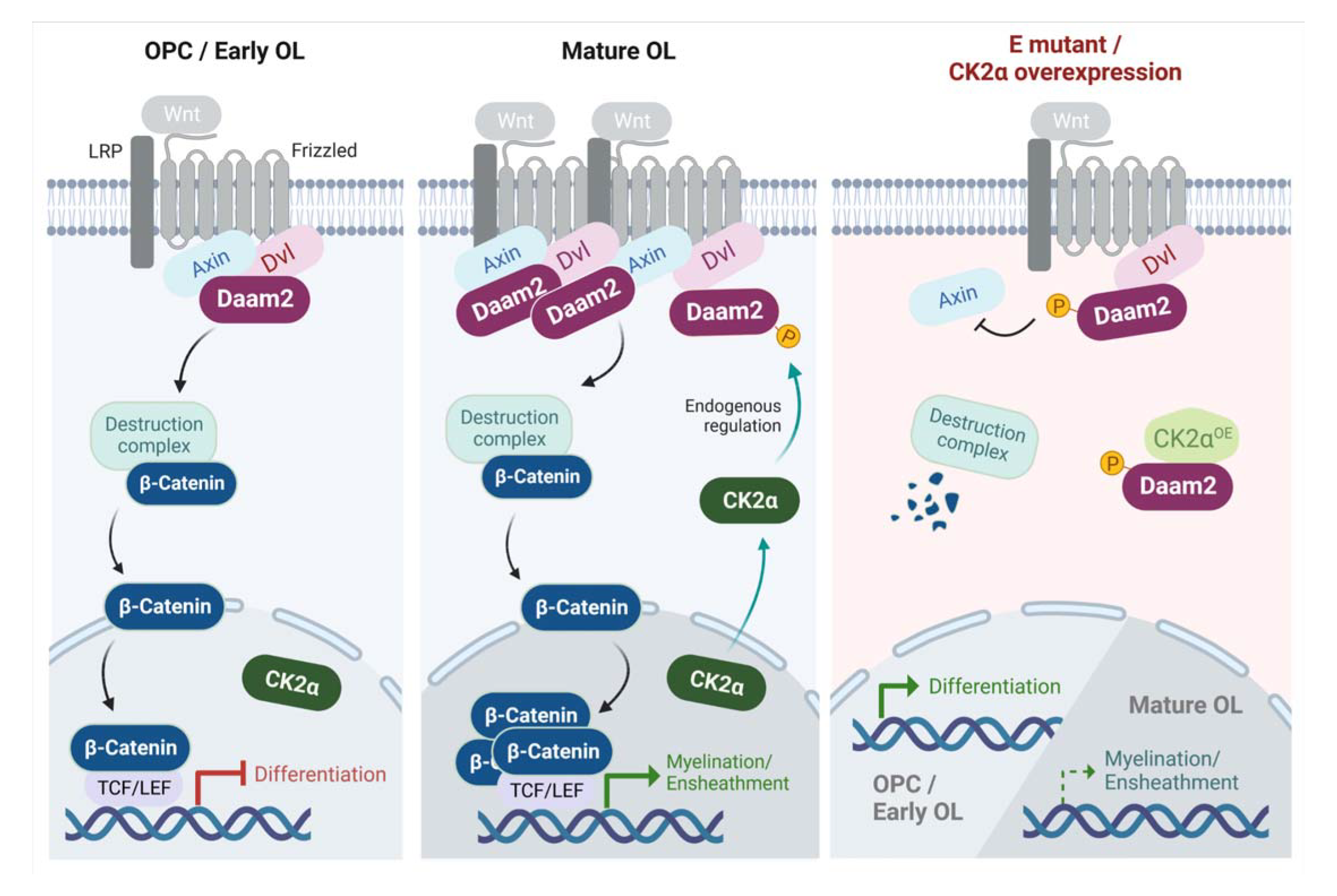
The molecular regulation of Wnt signaling in the OL lineage by Daam2 phosphorylation. Activity of Daam2-mediated Wnt signaling is low in OPCs and early OLs (left panel) but becomes upregulated in mature OLs (middle panel). Once the ligands trigger Wnt signaling, β-catenin will enter the nucleus where it inhibits and enhances expression of differentiation genes and myelination/ensheathment genes respectively. CK2α stays in the nucleus and does not interact with Daam2 in early OLs. In contrast, CK2α which translocates to cytosol in late OLs can phosphorylate Daam2 at S704/T705, followed by interference with Daam2-Axin2 interaction. Finally (Right panel), the E-mutation or CK2α overexpression weakens the Wnt signaling complex, thereby facilitating OL differentiation while negatively impacting ensheathment. Figure was created with BioRender.com.

## Discussion

In this study, we present a new regulatory pathway involved in the progression of OL lineage during brain development and white matter repair. Firstly, we demonstrated the effect of Daam2-mediated canonical Wnt signaling at specific developmental stages of the OL lineage. Secondly, we identified a novel post-translational regulation that involves CK2α, which interferes with the interaction between Daam2 and Axin2 in the Wnt signaling complex, and promotes OL differentiation through Daam2 phosphorylation. Our white matter injury models, mimicking HIE and MS, confirm that excessive Wnt activity significantly impedes developmental and regenerative myelination by inhibiting OL differentiation. However, targeting CK2α-mediated Daam2 phosphorylation can increase the pool of available OLs and promote white matter repair, even if the process of ensheathment might be slower after injury. Overall, our findings suggest that the CK2α-Daam2 pathway plays a crucial role in the regulation of OL biology and pathology associated with Wnt signaling.

The function of Wnt signaling in OL development has been investigated in various genetic animal models (7, 12). However, due to the limitations of *in vivo* models, the functional complexity and stage-specific roles of Wnt and its components are not fully understood. For instance, the destruction complex proteins, GSK3β, APC, and Axin2, as well as the β-catenin transcription partner Tcf7l2, are dispensable or participate in other pathways (32-35). Recently, Tcf7l2 was proposed to serve considered as a promoter for premyelinating OLs rather than an inhibitor (36). Furthermore, the genetic approach of combining OL-specific promoters (e.g., *NG2* vs *PLP*) with a *CreERT* system has not been used to compare early and late events in white matter development. Importantly, regulating β-catenin activation through axon 3 and 2-6 deletion may not recapitulate a nuclear translocation event in OL lineage, as compared to the conventional Wnt pathway in endothelial cells (37, 38). In this study, we show the stage-specific role of Wnt signaling in OL lineage progression. Expression of nuclear β-catenin levels suggest that higher endogenous Wnt activity in late stage of OLs maturation than in early OL development, which is well correlates with the Daam2 expression pattern in OL lineage progression. In addition, we prove Daam2 controls ligand-based Wnt activity in stage-specific OL development, which can be regulated by CK2α induced Daam2 phosphorylation *in vitro* and *in vivo*. Moreover, Daam2 phosphorylation regulates different population of Wnt-related genes in distinct OPC/OL clusters from our scRNA-seq analysis. Our findings illustrate the dynamic mechanistic basis of canonical Wnt signaling via Daam2 in early and late OL progression.

FH2, a conserved domain in the formin protein family, can bind to F-actin and nucleate actin filaments (39). However, little is known about its function after phosphorylation modification. Although FH2 phosphorylation may not affect nucleation activity (40), FH2 phosphorylated by CK2 disrupts the interaction between formin protein FHOD3 and ubiquitin-binding scaffold protein SQSTM1 (41). This raises the possibility that S704/T705 phosphorylation at the Daam2 FH2 domain weakens the binding to unknown partners (including Axin2), which are important for Wnt signalosome complex formation (15). On the other hand, our recent study shows that Daam2 interacts with the E3 ligases VHL and Nedd4, which reciprocally control OL development in the spinal cord (17). It is possible that other E3 ligases can be involved in Daam2-mediated Wnt signaling, since ubiquitination can also control protein interaction and signaling (42). Here, we report the novel role of phosphorylation modification at Daam2 in OL development, yet the further signaling cascade is needed for investigation.

Under normal conditions, CK2α-Daam2 interaction remains low in the cytosol compartment of OPCs and early OLs but it increases in late OLs. Because CK2 also enhances Wnt signaling through phosphorylating Dvl and β-catenin (20), this suggests a negative feedback for Wnt when CK2α translocated to cytosol. This raises an important question about the mechanism for CK2α shuttling between the nucleus and cytosol in OLs. CK2α may form different holoenzymes with other subunits (α‘ and β) in the nucleus and cytoplasm and display distinct catalytic activities and functions. Accordingly, the nuclear accumulation of β-catenin in APC-enriched mature OLs may result from cytosolic CK2α which phosphorylates β-catenin and releases it from APC and Axin2 in the destruction complex (7). On the other hand, in pathological conditions, Daam2 and Wnt signaling are upregulated to inhibit OL differentiation (9, 17), but CK2α is restricted to the nucleus in OPCs and early OLs and unable to interact with Daam2. To overcome this challenge, we manipulated the regulatory pathway by increasing cytosolic CK2α to successfully facilitate remyelination after injury. This result highlights the translational significance of pharmaceutical reagents capable of inducing cytosolic translocation of CK2α and Daam2 phosphorylation for demyelination diseases.

Future research should further investigate the roles of CK2 in OLs. For example, CK2 blockage with cx-4945 (Silmitasertib) is a therapeutic candidate for multiple sclerosis for its ability to disrupt T cell development (43, 44). Additionally, inhibition of CK2 activity was found to preserve OLs from AMPA-induced toxicity (45). However, we confirm an adverse effect of CK2 inhibitors on OL differentiation. CK2α overexpression also enhances OL differentiation *in vivo*. Similarly, CK2β plays an essential role in NSC proliferation and OPC production (22). CK2 kinase activity is also required for gene transcription of NG2 proteoglycan, which is a marker for OPCs and required for oligodendrogenesis (46). Given that CK2β can interact with OL-specific transcription factor Olig2 in a cell-free or immortalized cell line-based assay (22), CK2α may also functionally bind to Olig2 or other transcription factors during OL development (47). Together, CK2-targeting therapies could be exploited for OL survival in the acute phase of white matter diseases and for OL differentiation in the recovery stage.

## Acknowledgement

We thank Diego Cortes and Dr. Carlo D. Cristobal for technical assistance and also appreciate Dr. Etty (Tika) Benveniste for sharing reagents. This work was supported by grants from NIH/NINDS (R01NS110859 to H.K.L.), National Multiple Sclerosis Society (RG-1907-34551 to H.K.L.), and the Mark A. Wallace Endowment established by an anonymous donor (to H.K.L.), Electron microscopy, behavioral assays, and morphological analysis were supported in part by the Eunice Kennedy Shriver National Institute of Child Health & Human Development of the National Institutes of Health under award number P50HD103555 for the use of the BCM IDDRC Neurobehavior and Neurovisualization Cores. Mass spectrometry was supported by Dr. Sung Yun Jung and Mass Spectrometry Proteomics core at Baylor College of Medicine. scRNA-sequencing was partially supported by SCG core (S10OD023469, S10OD025240, P30EY002520, RP20054) and GARP core (1S10OD023469).

## Author contributions

C.-Y.W., and H.K.L. conceived the project and designed the experiments. C.-Y.W. performed the all the experiments. K.I.K. performed animal behavior tests. Z.Z. performed electron microscopy. H.J.B. provided reagents and expertise. C.-Y.W. and H.K.L. wrote the manuscript.

## Competing interests

The authors declare no competing interests.

## Materials and Methods

### Materials

All experimental materials are included in **Table S2**.

### Mice

To obtain Daam2 phospho-mimetic (E) mutation mouse line, the point mutation was generated by CRISPR/Cas9-base genomic editing. The mouse line was created after ES cell microinjection in Genetically Engineered Rodent Models Core at the Baylor College of Medicine. Genetic mutation was confirmed by PCR using specific primers. Flag-Daam2 knock-in mice have been used in our previous study (48). All procedures were approved by the Institutional Animal Care and Use Committee (IACUC) at Baylor College of Medicine and conform to the US Public Health Service Policy on Humane Care and Use of Laboratory Animals.

### Primary OPC/OL culture

Cortical tissues were collected from the mouse embryos at E14.5, and the neural stem cells (NSCs) were maintained as neurospheres in DMEM-F12 plus B27, N2 supplements, 20 ng/ml EGF and 20 ng/ml FGF for 4 days. For OPC induction, NSCs were dissociated with Accutase and plated onto poly-D-lysine coated dishes and coverslips in OPC medium (DMEM-F12 plus B27, 20 ng/ml FGF and 20 ng/ml PDGF-AA) for 2 days. To generate OLs, OPCs were then differentiated in OL medium, consist of DMEM, 1% GlutaMax, 1 mM sodium pyruvate, 0.1% BSA, 50 μg/ml apotransferrin, 5 μg/ml insulin, 30 nM sodium selenite, 10nM D-biotin, 10 nM hydrocortisone, 5 μg/ml NAC, 10 ng/ml T3, and 10 μg/ml CNTF for 2 days (early stage) and 4 days (late stage). For gene transduction, OPCs were transfected with Flag-tagged Daam2 and specific plasmids by iMfectin transfection agents for 16 hours prior to differentiation for further experiments.

### Single cell RNA sequencing (scRNA-seq)

P21 mouse cortical tissues containing corpus callosum (3 animals combined per group) were dissociated with papain at 37 degree for 30 min. Cell solution was prepared using debris and dead cell removal kit with MS separation columns. Single cell gene expression library was prepared according to Chromium Single Cell Gene Expression 3v3.1 kit (10x Genomics) by Single Cell Genomic Core (SCG) at the Baylor College of Medicine. After quality control of cDNA libraries, primary data were processed with Illumina Next Generation Sequencing by Genomic & RNA Profiling Core (GARP) at the Baylor College of Medicine. The scRNA-seq data were analyzed with Cell Ranger Count v6.1.2 followed by R-studio. Cells with more than 7% mitochondrial RNA or less than 1000 total RNA counts were excluded from data.

### In vitro CK2 kinase assay

To determine whether CK2α directly phosphorylates Daam2, purified Daam2 proteins and short synthetic peptides were analyzed using ADP-Glo CK2α kinase assay. The peptides and casein (positive control) were incubated with recombinant CK2α and ATP in reaction buffer at 37 degree for 60 min. The levels of ADP generated from phosphorylation reactions were then determined by luminescence.

### Immunofluorescence

Tissues collected from mice were fixed in 4% paraformaldehyde (PFA) and dehydrated with 30% sucrose. Tissues were embedded in OCT for frozen section and cut at 20 μm-thickness. Tissue sections were permeabilized with PBS containing 1% triton X-100 (PBST) and blocked using 1% BSA. Primary antibodies in 0.3% PBST were applied onto tissue sections at 4 degree overnight. Subsequently, tissue sections were incubated with fluorophore conjugated secondary antibodies for 60 min followed by nucleus counterstaining with DAPI. Otherwise, cells on coverslips were fixed with 4% PFA and treated with 0.1% PBST. Coverslips were sequentially incubated with primary and secondary antibodies in PBS, followed by DAPI. Samples were mounted with VectaShield anti-fade mounting medium. Stained tissues and cells were imaged under a Zeiss Imager.M2m microscope equipped with ApoTome.2, AxioCam 506 mono, and AxioCam MRc.

### In situ hybridization

Detection of gene transcripts in the brain tissues were performed as previous described (17). RNA probes of MBP, PLP, PDGFRα conjugated with DIG were generated in-house for in situ hybridization. Briefly, tissue sections were post-fixed in 4% PFA and treated with proteinase K and acetyl anhydride solution. After incubation with RNA probes at 65 degree overnight, tissue sections were washed with SSC buffer and the probes were detected using DIG antibody conjugated with alkaline phosphatase. The RNA signals were visualized by NBT/BCIP substrates.

### Quantitative real-time PCR (Q-PCR)

Q-PCR analysis was performed as described previously (17). Briefly, Total RNA from corpus callosum tissue of P21 mouse brains was extracted using TRIzol. Q-PCR was conducted in Bio-Rad real-time PCR systems using the amfiSure PCR master mix following the manual. Relative mRNA expression level was determined by the threshold cycle (Ct) value of each PCR product and normalized to that of GAPDH.

### Immunoprecipitation and western blot

Tissues or cells were homogenized in SDS-free RIPA buffer plus protease and phosphatase inhibitors and then incubated with primary antibodies and protein A/G beads at 4 degree overnight. The beads containing IP complex after washed with RIPA buffer or tissue/cell lysates were boiled with loading buffer. Proteins were separated by 8-12% SDS-PAGE and transferred to nitrocellulose membranes. The membranes were blotted with primary antibodies and HRP-conjugated secondary antibodies. Protein signals were visualized by luminescence using ECL.

### Electron microscopy

Tissue preparation and processing followed our previous study (18). Mice were perfused with PBS and fixative solution (1% glutaraldehyde, 4% paraformaldehyde, 0.1 m sodium cacodylate). Regions of interest located in the corpus callosum were dissected at 1 mm^3^ blocks and post-fixed at 4 degree for additional 1 hour. Lipid content was fixed with 0.1M cacodylate solution containing 1% osmium tetroxide, 1.5% KFeCN at 4 degree for 1 hour. Next, the tissue blocks were washed with 0.1M cacodylate and fixed again in 2% glutaraldehyde at 4 degree overnight followed by washing and dehydration steps. Tissue blocks were then embedded in resin and cured for 5 days. The tissue blocks were randomly numbered before sectioning and underwent single-blinded analysis by electron microscopy.

### Neonatal hypoxic injury and 3-chamber test

Based on the protocol from our previous study (16), P3 mouse pups and nursing mother were placed into a chamber, and the oxygen level was adjusted and maintained at 10.5% by nitrogen. After 8-day incubation, mice were returned to normoxic environment for 7 days. Challenged mice were harvested at P11 to confirm hypomyelination and at P18 to evaluate myelination level. For the 3-chamber sociability test, 2-month-old mice were randomly number-tagged before single-blinded analysis. Mice were placed in a middle chamber which connects to chambers on the left and right side, and mice were allowed to explore the 3 chambers for 10 min in each trial. The location and movement of the mice were recorded by a camera. In first trial for habituation, an object was placed in the left and right chambers respectively. In second trial, a new object was put in the left chamber and a stranger mouse in the right chamber respectively. In third trial, the object from second trial was replaced by a new stranger mouse, and the mouse in the right become a familiar mouse.

### Lysolecithin-induced demyelination

As previously described (16, 17), mice were anesthetized and were placed on a stereotaxic device. After skull was exposed, a hole was made at 1-mm anterior and 1-mm right to the bregma. 1 μl PBS containing 2% lysolecithin and ∼10^9^ U AAVs were injected into the brain at the depth of 1.5 mm. The brain containing lesion at corpus callosum was collected at 14 days post injection.

### Image analysis, quantification, and statistics

Protein signals from western blot (non-blinded), cell number counting, signal intensity, area, and sholl analysis from immunofluorescence (single-blinded) were measured using ImageJ software with related plug-ins. For electron microscopy (single-blinded), g-ratio measurement, radius of axons with or without myelin layers were measured manually using ImageJ software. Data were analyzed and the result plots were generated using Prism 6 software. For data comparison, student t-test and one-way ANOVA were used when only one variable factor between groups. Two-way ANOVA with Sidak’s multiple comparison test was used for two variables among groups.

### Data, Materials, and Software Availability

Any requirement for the data and scripts in this study, additional information, or reanalyzing the data reported in this study is available from the correspondence upon request.

## Notes

### Competing Interest Statement

The authors have declared no competing interest.

## References

1. Philips T & Rothstein JD (2017) Oligodendroglia: metabolic supporters of neurons. J Clin Invest 127(9):3271–3280.

2. Ziemka-Nalecz M, et al. (2018) Impact of neonatal hypoxia-ischaemia on oligodendrocyte survival, maturation and myelinating potential. J Cell Mol Med 22(1):207–222.

3. Dulamea AO (2017) Role of Oligodendrocyte Dysfunction in Demyelination, Remyelination and Neurodegeneration in Multiple Sclerosis. Adv Exp Med Biol 958:91–127.

4. Buser JR, et al. (2012) Arrested preoligodendrocyte maturation contributes to myelination failure in premature infants. Ann Neurol 71(1):93–109.

5. Kuhlmann T, et al. (2008) Differentiation block of oligodendroglial progenitor cells as a cause for remyelination failure in chronic multiple sclerosis. Brain 131(Pt 7):1749–1758.

6. Patel JR & Klein RS (2011) Mediators of oligodendrocyte differentiation during remyelination. FEBS Lett 585(23):3730–3737.

7. Wheeler NA & Fuss B (2016) Extracellular cues influencing oligodendrocyte differentiation and (re)myelination. Exp Neurol 283(Pt B):512–530.

8. Lock C, et al. (2002) Gene-microarray analysis of multiple sclerosis lesions yields new targets validated in autoimmune encephalomyelitis. Nat Med 8(5):500–508.

9. Han MH, et al. (2008) Proteomic analysis of active multiple sclerosis lesions reveals therapeutic targets. Nature 451(7182):1076–1081.

10. Xie C, Li Z, Zhang GX, & Guan Y (2014) Wnt signaling in remyelination in multiple sclerosis: friend or foe? Mol Neurobiol 49(3):1117–1125.

11. Fancy SP, et al. (2009) Dysregulation of the Wnt pathway inhibits timely myelination and remyelination in the mammalian CNS. Genes Dev 23(13):1571–1585.

12. Guo F, et al. (2015) Canonical Wnt signaling in the oligodendroglial lineage--puzzles remain. Glia 63(10):1671–1693.

13. Ghelman J, et al. (2021) SKAP2 as a new regulator of oligodendroglial migration and myelin sheath formation. Glia 69(11):2699–2716.

14. Kawabata Galbraith K & Kengaku M (2019) Multiple roles of the actin and microtubule-regulating formins in the developing brain. Neurosci Res 138:59–69.

15. Lee HK & Deneen B (2012) Daam2 is required for dorsal patterning via modulation of canonical Wnt signaling in the developing spinal cord. Dev Cell 22(1):183–196.

16. Lee HK, et al. (2015) Daam2-PIP5K is a regulatory pathway for Wnt signaling and therapeutic target for remyelination in the CNS. Neuron 85(6):1227–1243.

17. Ding X, et al. (2020) The Daam2-VHL-Nedd4 axis governs developmental and regenerative oligodendrocyte differentiation. Genes Dev 34(17-18):1177–1189.

18. Cristobal CD, et al. (2022) Daam2 Regulates Myelin Structure and the Oligodendrocyte Actin Cytoskeleton through Rac1 and Gelsolin. J Neurosci 42(9):1679–1691.

19. Clevers H & Nusse R (2012) Wnt/beta-catenin signaling and disease. Cell 149(6):1192-1205.

20. Seldin DC, et al. (2005) CK2 as a positive regulator of Wnt signalling and tumourigenesis. Mol Cell Biochem 274(1-2):63–67.

21. Dominguez I, Sonenshein GE, & Seldin DC (2009) Protein kinase CK2 in health and disease: CK2 and its role in Wnt and NF-kappaB signaling: linking development and cancer. Cell Mol Life Sci 66(11-12):1850–1857.

22. Huillard E, et al. (2010) Disruption of CK2beta in embryonic neural stem cells compromises proliferation and oligodendrogenesis in the mouse telencephalon. Mol Cell Biol 30(11):2737–2749.

23. Castello J, Ragnauth A, Friedman E, & Rebholz H (2017) CK2-An Emerging Target for Neurological and Psychiatric Disorders. Pharmaceuticals (Basel) 10(1).

24. Wisniewski JR, Nagaraj N, Zougman A, Gnad F, & Mann M (2010) Brain phosphoproteome obtained by a FASP-based method reveals plasma membrane protein topology. J Proteome Res 9(6):3280–3289.

25. Xia Q, et al. (2008) Phosphoproteomic analysis of human brain by calcium phosphate precipitation and mass spectrometry. J Proteome Res 7(7):2845–2851.

26. Blom N, Sicheritz-Ponten T, Gupta R, Gammeltoft S, & Brunak S (2004) Prediction of post-translational glycosylation and phosphorylation of proteins from the amino acid sequence. Proteomics 4(6):1633–1649.

27. Millar LJ, Shi L, Hoerder-Suabedissen A, & Molnar Z (2017) Neonatal Hypoxia Ischaemia: Mechanisms, Models, and Therapeutic Challenges. Front Cell Neurosci 11:78.

28. Wang Y & Olson IR (2018) The Original Social Network: White Matter and Social Cognition. Trends Cogn Sci 22(6):504–516.

29. Giannopoulou I, Pagida MA, Briana DD, & Panayotacopoulou MT (2018) Perinatal hypoxia as a risk factor for psychopathology later in life: the role of dopamine and neurotrophins. Hormones (Athens) 17(1):25–32.

30. van Handel M, Swaab H, de Vries LS, & Jongmans MJ (2010) Behavioral outcome in children with a history of neonatal encephalopathy following perinatal asphyxia. J Pediatr Psychol 35(3):286–295.

31. Blakemore WF & Franklin RJ (2008) Remyelination in experimental models of toxin-induced demyelination. Curr Top Microbiol Immunol 318:193–212.

32. Azim K & Butt AM (2011) GSK3beta negatively regulates oligodendrocyte differentiation and myelination in vivo. Glia 59(4):540–553.

33. Lang J, et al. (2013) Adenomatous polyposis coli regulates oligodendroglial development. J Neurosci 33(7):3113–3130.

34. Dai ZM, et al. (2014) Stage-specific regulation of oligodendrocyte development by Wnt/beta-catenin signaling. J Neurosci 34(25):8467–8473.

35. Hammond E, et al. (2015) The Wnt effector transcription factor 7-like 2 positively regulates oligodendrocyte differentiation in a manner independent of Wnt/beta-catenin signaling. J Neurosci 35(12):5007–5022.

36. Guo F & Wang Y (2023) TCF7l2, a nuclear marker that labels premyelinating oligodendrocytes and promotes oligodendroglial lineage progression. Glia 71(2):143–154.

37. Ye F, et al. (2009) HDAC1 and HDAC2 regulate oligodendrocyte differentiation by disrupting the beta-catenin-TCF interaction. Nat Neurosci 12(7):829–838.

38. Yu W, McDonnell K, Taketo MM, & Bai CB (2008) Wnt signaling determines ventral spinal cord cell fates in a time-dependent manner. Development 135(22):3687–3696.

39. Schonichen A & Geyer M (2010) Fifteen formins for an actin filament: a molecular view on the regulation of human formins. Biochim Biophys Acta 1803(2):152–163.

40. Moseley JB & Goode BL (2005) Differential activities and regulation of Saccharomyces cerevisiae formin proteins Bni1 and Bnr1 by Bud6. J Biol Chem 280(30):28023–28033.

41. Iskratsch T, et al. (2010) Formin follows function: a muscle-specific isoform of FHOD3 is regulated by CK2 phosphorylation and promotes myofibril maintenance. J Cell Biol 191(6):1159–1172.

42. Mukhopadhyay D & Riezman H (2007) Proteasome-independent functions of ubiquitin in endocytosis and signaling. Science 315(5809):201–205.

43. Crunkhorn S (2016) Autoimmune disease: CK2 blockade ameliorates EAE. Nat Rev Drug Discov 15(11):750.

44. Ulges A, et al. (2016) Protein kinase CK2 governs the molecular decision between encephalitogenic TH17 cell and Treg cell development. Proc Natl Acad Sci U S A 113(36):10145–10150.

45. Canedo-Antelo M, et al. (2018) Inhibition of Casein Kinase 2 Protects Oligodendrocytes From Excitotoxicity by Attenuating JNK/p53 Signaling Cascade. Front Mol Neurosci 11:333.

46. Schmitt BM, et al. (2020) Protein Kinase CK2 Regulates Nerve/Glial Antigen (NG)2-Mediated Angiogenic Activity of Human Pericytes. Cells 9(6):1546.

47. Zhou J, et al. (2017) A Sequentially Priming Phosphorylation Cascade Activates the Gliomagenic Transcription Factor Olig2. Cell Rep 18(13):3167–3177.

48. Jo J, et al. (2021) Regional heterogeneity of astrocyte morphogenesis dictated by the formin protein, Daam2, modifies circuit function. EMBO Rep 22(12):e53200.

