## Supporting information for "CK2α-dependent regulation of Wnt activity governs white matter development and repair"

**Figure S1**

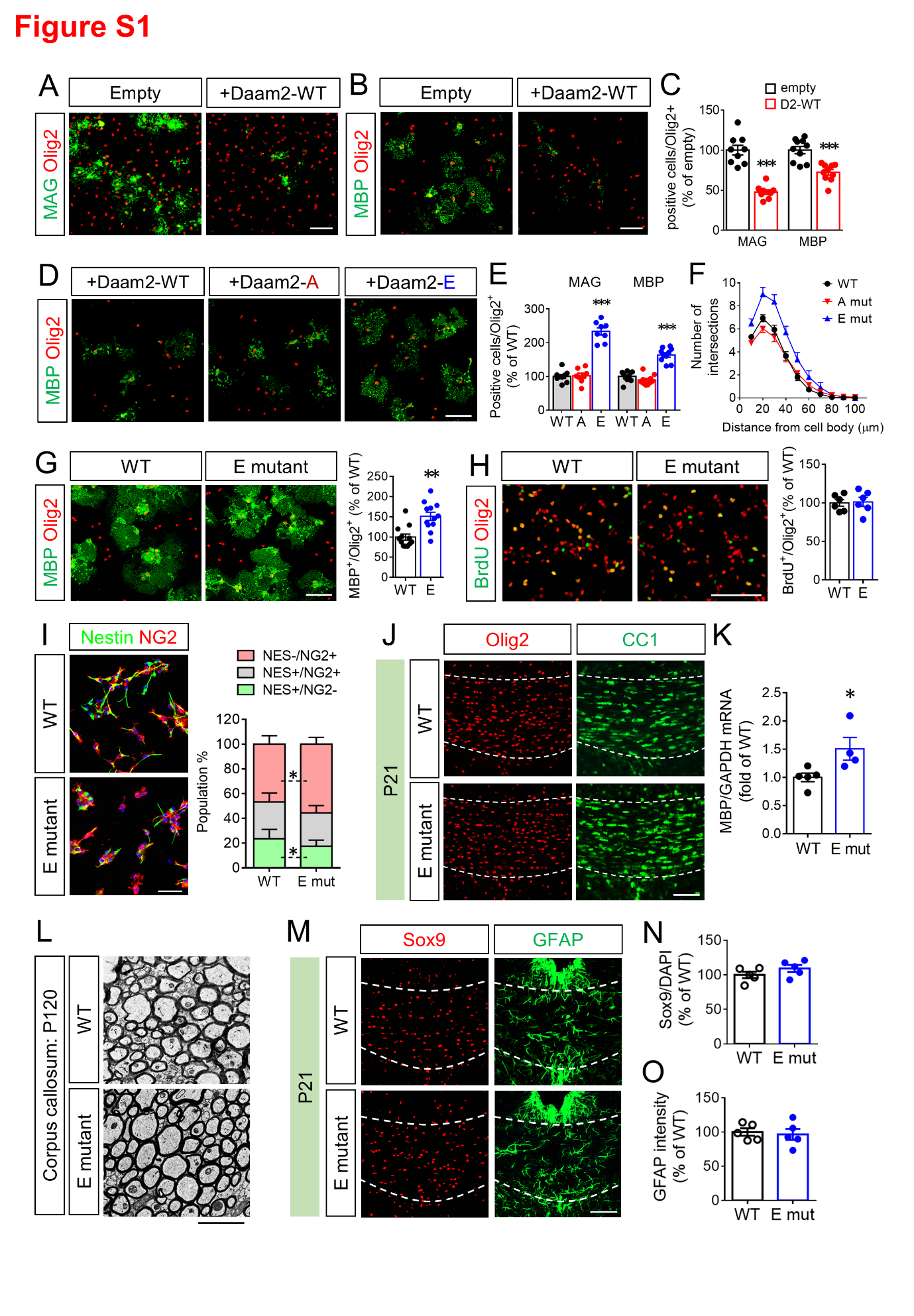

**Figure S1. Phospho-mimetic mutation of Daam2 is beneficial for OL differentiation.** Primary OPCs were transfected with Flag-Daam2 and differentiated for 2 days (A) and 4 days (B, D). (C, E) In vitro differentiation was assessed by immunofluorescence for MAG and MBP. (F) The number of cell processes and branches extend from a cell body at different distances were calculated using sholl analysis for process complexity. (G) OPCs from the E mutant were differentiated for 4 days followed by immunofluorescence for MBP. (H) WT and the E mutant OPCs were treated with BrdU for 6 hrs before immunofluorescence to detect proliferating cells with BrdU incorporation. (I) WT and the E-mutant NSCs were differentiated into OPCs for 16 hrs followed by immunofluorescence for the markers for NSCs (nestin) and OPCs (NG2). P21 corpus callosum from WT and the E mutant were analyzed by immunofluorescence (J, M) and by Q-PCR (K). (L) The myelin structure in the corpus callosum from WT and the E-mutant mice at P120 were subjected to electron microscopy. The number of Sox9+ cells was counted (N), and the immunoreactivity of GFAP was measured (O). Data from at least 3 independent experiments or animals for each group were presented as mean ± SEM. *P < 0.05, ***P < 0.001 versus empty in C, versus WT in E, K. Scale bar, 100 μm in all except L; 2 μm in L.

**Figure S2**

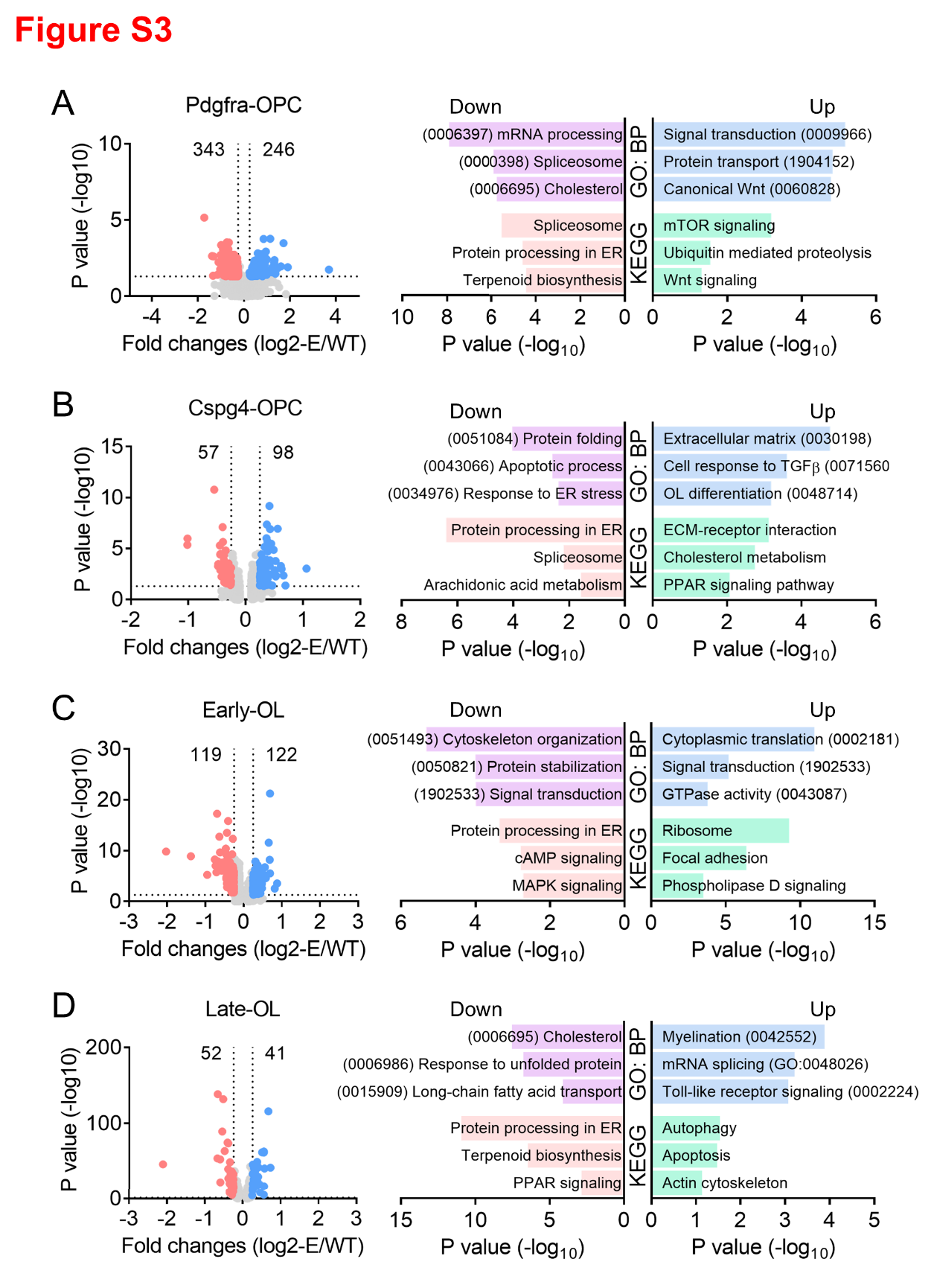

**Figure S2. Differentially expressed genes (DEGs) in 4 OPC/OL clusters between WT and the E mutant brains.** DEGs with fold change (Log_2_) > 0.25 and P value < 0.05 and their numbers are shown in the volcano plots. Gene ontology and KEGG pathway analysis were performed, and the Top 3 candidates related to OL biological function are listed.

**Figure S3**

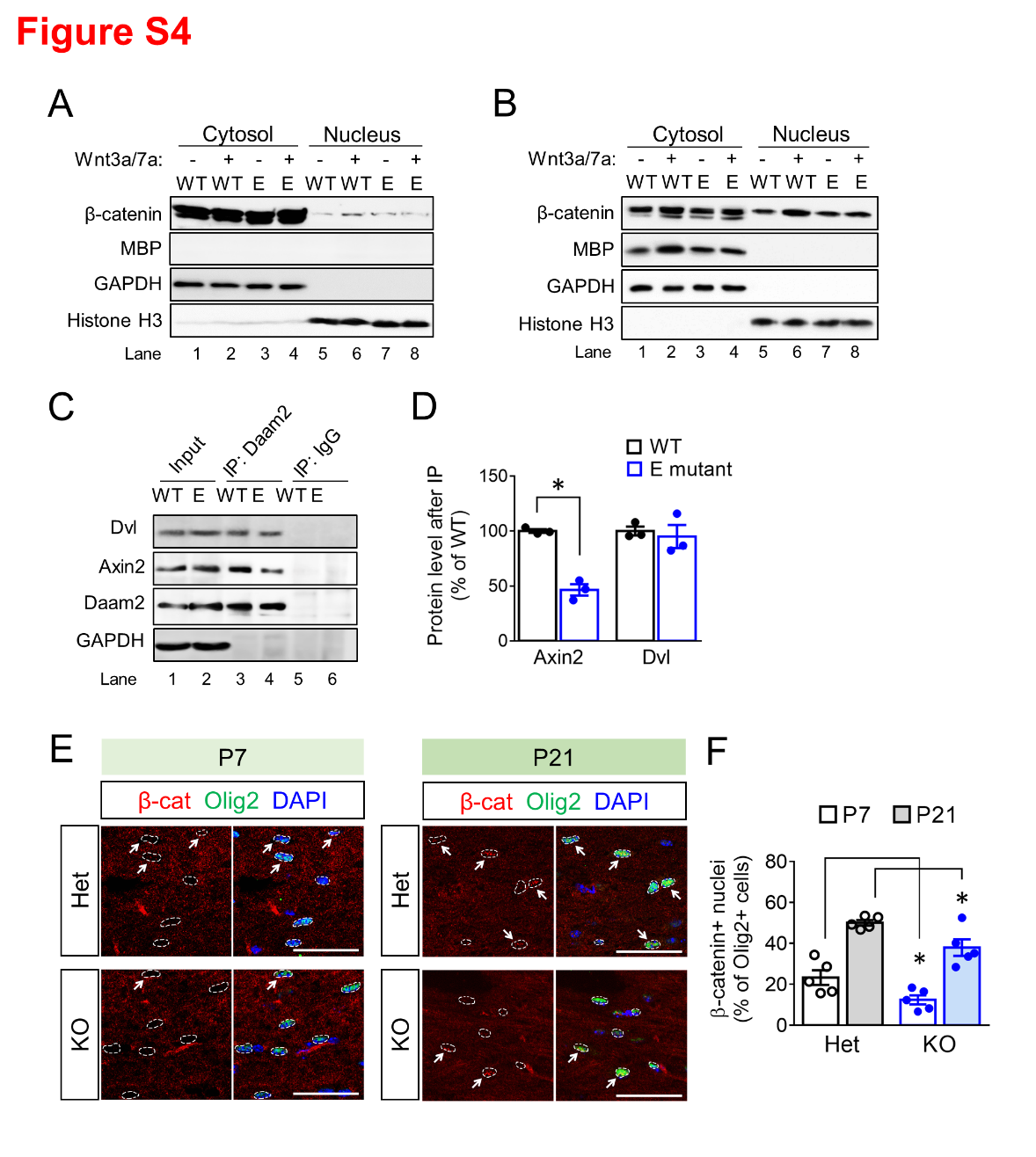

**Figure S3. Daam2-E mutation blocks ligand-based Wnt activation in OLs.** (A-B) Protein levels in the cytosol and nucleus fractions of early OLs (A) and late OLs (B) were analyzed by western blot. GAPDH serves as a loading control for cytosol fractions, and Histone H3 for nucleus fractions.

**Figure S4**

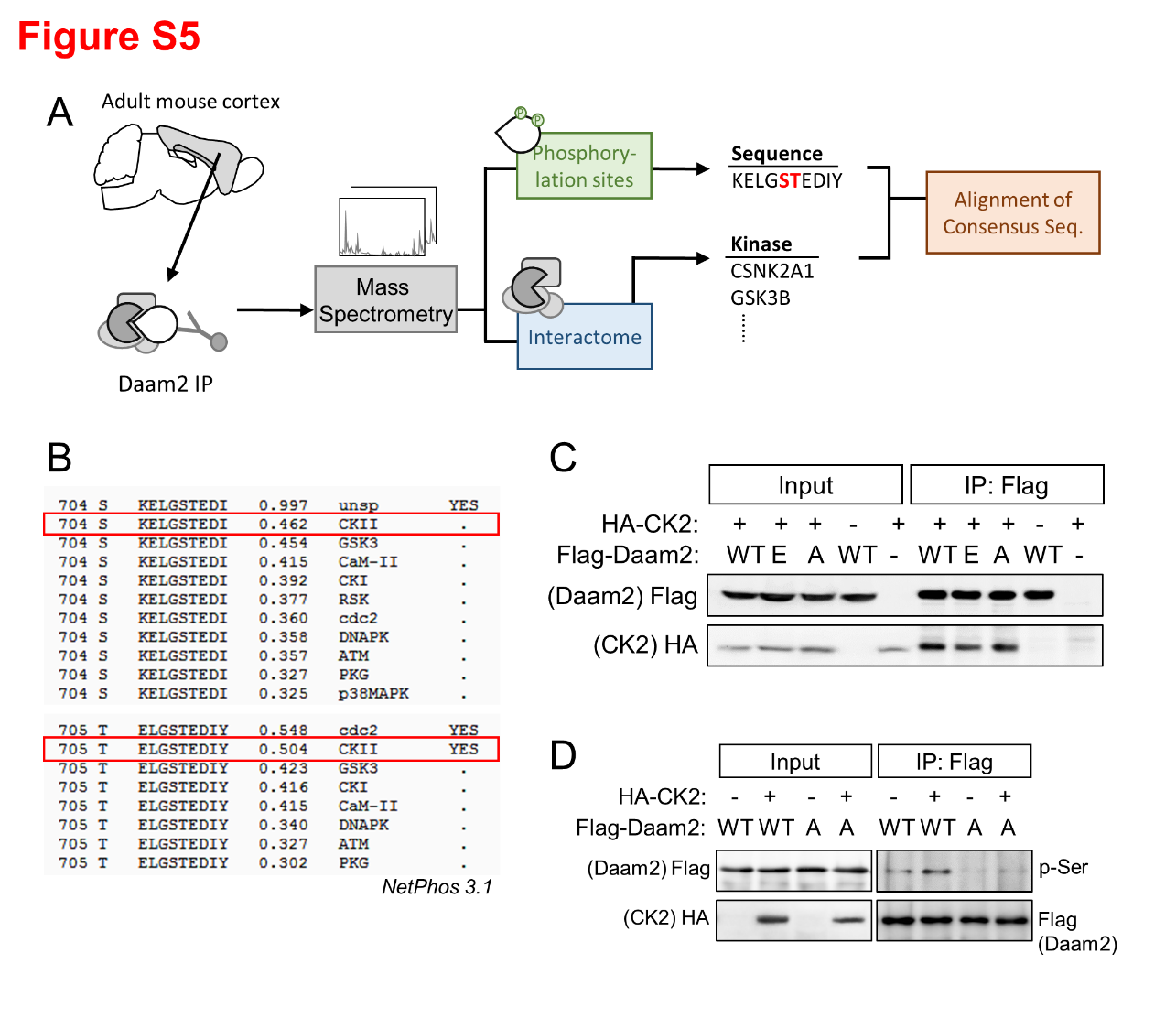

**Figure S4. CK2α is a candidate to phosphorylate Daam2 at S704/T705.** (A) Experiment design for identifying candidate kinases phosphorylating Daam2. (B) The phosphorylation motif at S704/T705 was aligned in NetPhos3.1 database for kinase prediction. (C) After transfecting primary OPCs with Flag-Daam2 and HA-CK2α, HA-CK2α was co-immunoprecipitated by anti-Flag. (D) Flag-Daam2 (WT and A-mut) and HA-CK2α were transfected into OPCs followed by differentiation for 2 days. Phospho-serine (p-Ser) levels on Flag-Daam2 were analyzed by western blot after Flag immunoprecipitation.

**Figure S5**

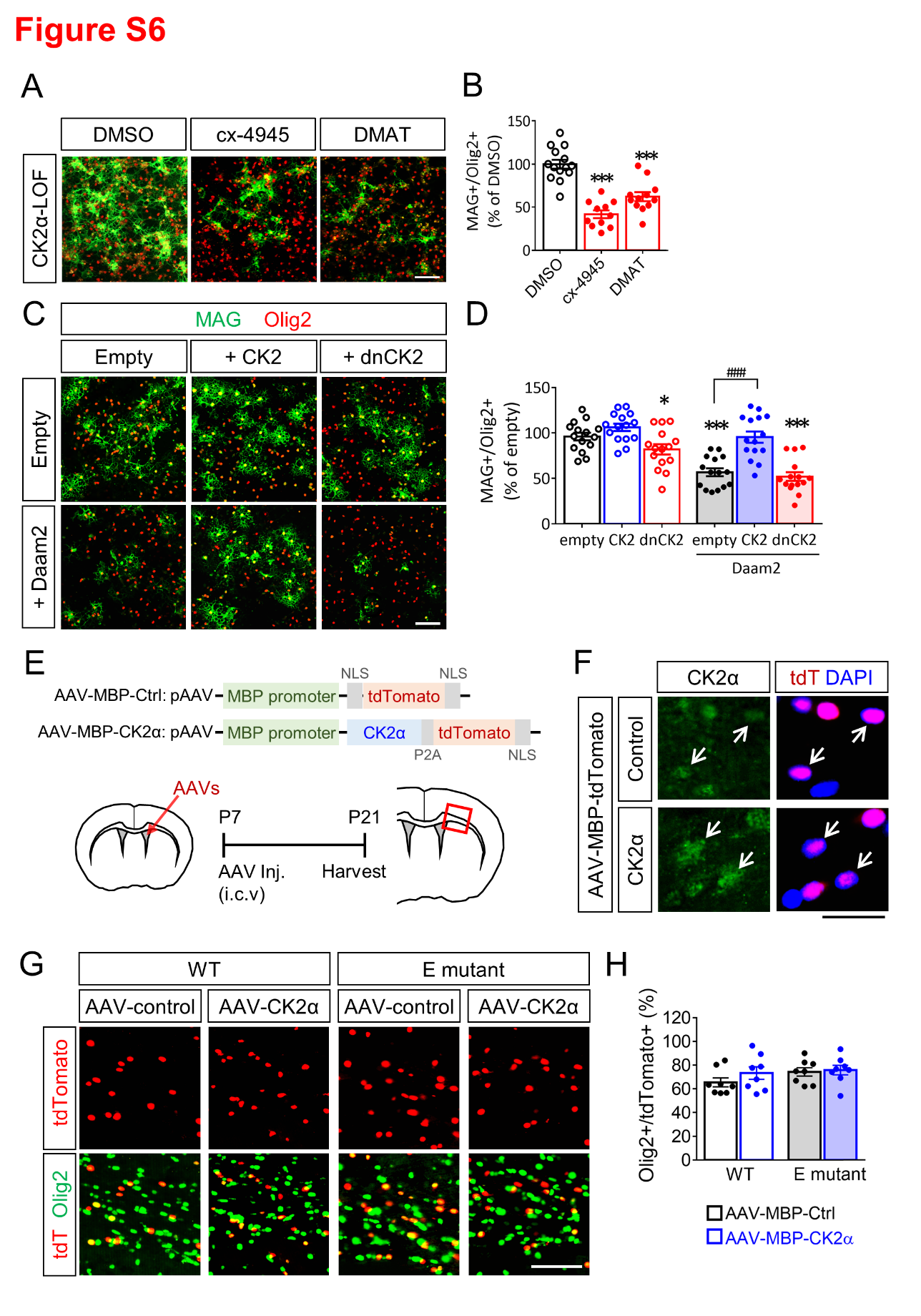

**Figure S5. CK2α kinase activity is required for OL differentiation.** (A-B) Primary OPCs were treated CK2 inhibitors and subjected to in vitro differentiation. (D) In vitro differentiation of OPCs transfected with Daam2 and CK2α (or dominant negative CK2α) were evaluated. (F) A diagram shows AAV constructs and the injection of AAVs into the brain. (G) The overexpression of CK2α in tdtomato^+^ cells in P21 corpus callosum injected with AAV-MBP-CK2α. (H-I) The brains injected with AAVs were assessed by immunofluorescence at P21. The number of Olig2^+^ and tdtomato^+^ cell in the corpus callosum were counted. Data from at least 3 independent experiments or animals for each group were presented as mean ± SEM. *P < 0.05, ***P < 0.001 versus DMSO in B, versus empty in E; ^###^P <0.001 versus Daam2 in E. Scale bar, 100 μm in A, D; 50 μm in G; 100 μm in H.

**Figure S6**

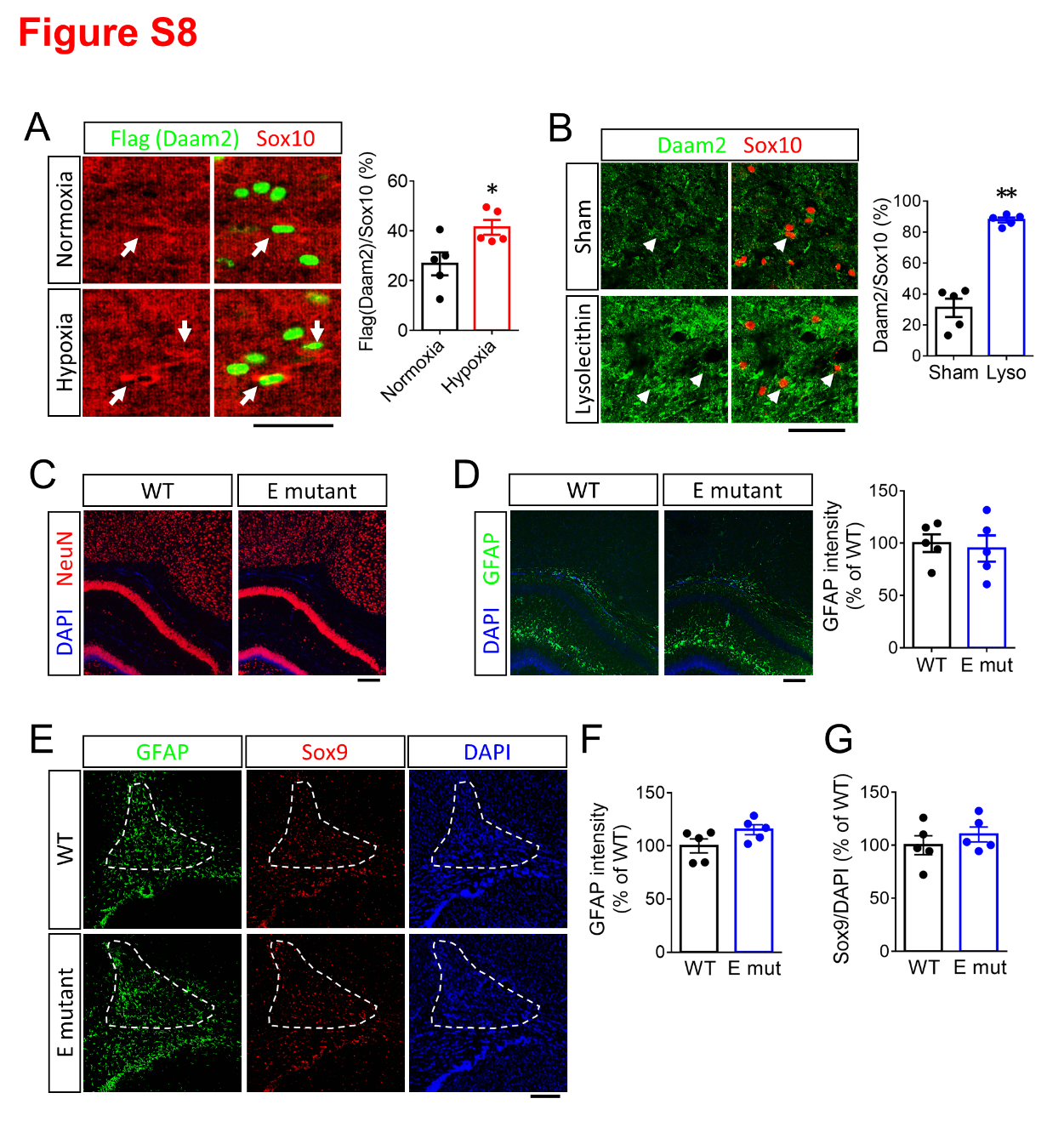

**Figure S6. Daam2 is upregulated after white matter injury.** P11 Flag-Daam2 knock-in brains after postnatal hypoxic injury (A) and the adult brains at 14 days after lysolecithin injection (B) were subjected to immunofluorescence. Intensity of Daam2 (or Flag) in Sox10^+^ cells in the corpus callosum was measured. (C-D) NeuN^+^ neurons and GFAP^+^ astrocytes were assessed in P18 brains after postnatal hypoxic injury. (E-G) GFAP^+^ and Sox9^+^ astrocytes were assessed at 14 days after lysolecithin injection. Data from at least 5 independent experiments or animals for each group were presented as mean ± SEM. *P < 0.05, **P < 0.01 versus normoxia in A, versus sham in B. Scale bar, 50 μm in A, 100 μm in B, 200 μm in C-E.
